# Transcriptomic and epigenetic assessment of ageing male skin identifies disruption of Ca^2+^ homeostasis; exacerbated by smoking and UV exposure

**DOI:** 10.1101/2022.08.16.504102

**Authors:** Louise I. Pease, James Wordsworth, Daryl Shanley

## Abstract

Skin ageing has been widely associated with the formation and presence of increasing quantities of senescent cells, the presence of which are thought to reduce cell renewal. This study aimed to identify key factors influencing fibroblast and skin aging in European males using RNA-seq data. Key differences in study designs included known sources of biological differences (sex, age, ethnicity), experimental differences, and environmental factors known to accelerate skin ageing (smoking, UV exposure) as well as study specific batch effects which complicated the analysis. To overcome these complications samples were stratified by these factors and differential expression assessed using Salmon and CuffDiff. Functional enrichment and consistency across studies, stratification’s and tools identified age related alterations in the transcriptomes of fibroblasts and skin. Functional enrichment of results identified alterations in protein targeting to membranes and the ER, and altered calcium homeostasis in aged fibroblasts. Extension to skin controlled for differences in fibroblast culturing methods confirming transient age related alterations in intracellular calcium homeostasis. In middle aged males (40-65) increased keratinisation, skin, epithelial and epidermal development was seen in conjunction with alterations to ER Ca^2+^ uptake, leading to the identification of related processes including; an unfolded protein response, altered metabolism, increased MMP expression, and altered Ca^2+^ handling, which were further exacerbated by UV-exposure. Interestingly the genes and processes subsequently decreased in old males (> 65), which exhibited signs of increased senescence. Extension to Illumina 450k array data from ageing skin uncovered evidence of epigenetic regulation; genes and isoforms with overlapping differentially methylated CpGs were differentially expressed. Smoking led to additional enrichment of genes relating to tissue development, cell adhesion, vasculature development, peptide cross-linking, calcium homeostasis, cancer and senescence. The results consistently identified alterations in ER and golgi Ca^2+^ uptake, which disrupt intracellular and extracellular calcium gradients that regulate metabolic and differentiation signalling in skin and fibroblasts, leading to age related declines skin structure and function. Interestingly many diseases and infections with overlapping molecular consequences, (ER Ca^2+^ stress, reduced protein targeting to membranes) including COVID-19 are identified by the analysis, suggesting that COVID-19 infection compounds pre-existing cellular stresses in aged males, which could help explain higher COVID-19 mortality rates in aged males, as well as highlighting potential ways to reduce them.

## Introduction

Skin is one of the most complicated organs in the human body, it is composed of three layers; The epidermis, dermis and subcutaneous tissue. Intrinsic ageing encompasses the spontaneous appearance of fine wrinkles and thinning of the epidermis with advancing age^1, 2^; influenced by ethnicity, anatomical variation, and hormones^3^, which differ by age and sex^4–6^. Extrinsic ageing encompasses accelerated changes in skin structure and function, evidenced by deep wrinkles, skin laxity, and hyperpigmentation, which can be brought about by chronic UV-exposure which alters the concentrations and localisation of adhesion proteins^1, 3–5^, pollutant exposure, excessive alcohol consumption, and smoking^1, 3, 4, 6–8^. The rate of ageing can be affected by alterations in the vascular / glandular network, and hydration^1, 7^. Wrinkling and sagging are caused by decreases in growth factors, collagen, and abnormal accumulation of elastin; flattening and thinning of the dermal epidermal junction^3, 7, 9^.

Male skin has more oil, sebum, lower trans-epidermal water loss and collagen and is 10-20 per cent thicker; however, collagen which decreases from the age of 20 years in males remains relatively constant until the age of 50 in females^6^. Females develop more dermatoses, thought to relate to sex hormone differences, infleunced by behavioral factors, as well as ethnicity; age related alterations in skin thickness, thought to relate to loss of dermal collagen, and loss of hydroxyl proline have been attributed to menopause and subsequent hormonal changes^6^. Overall skin differs depending on age, location on the body, geographical region, ethnicity and sex, which influences factors such as collagen content, oiliness, hydration, coarseness and elasticity^6^. For these reasons we limited this study to males to mitigate all of the aforementioned potential confounding factors (oil, collagen, hydroxyl proline content, sex hormones, and skin thickness) that are both different in males and females, and known to be affected by the ageing process.

### Previous studies

Transcriptomic analysis of cultured human dermal fibroblasts has been used alongside machine learning algorithms to predict the age of donors by Fleischer et al. (2018). Their data set consisted of 133 fibroblasts aged 0 - 89 and 10 from patients with Hutchinson-Gilford Progeria Syndrome (HGPS) between 2 and 8 years of age. HGPS is associated with premature ageing, alopecia and loss of subcutaneous fat, atherosclerosis, myocardial infarction and strokes, thought to be caused by mutations in the LMNA gene which codes for scaffolding nuclear laminins with roles in nuclear stability, chromatin structure and gene expression^10, 11^. The combined machine learning methods were accurate in predicting the age of donors, correctly identifying the HGPS samples and predicting HGPS patients were 9 to 10 years older than their chronological age, and age matched controls^11^. Kaisers et al. (2017) studied the impacts of age, gender and UV-exposure on gene expression in short term cultured fibroblasts^9^. In their study donors were grouped by gender and age (young 19-25 yrs, middle 36-45 yrs, Old 60-66 yrs) and each donor provided a UV-exposed and non UV-exposed sample. Histological analysis of samples identified age dependent loss of rete ridges, thinning of the epidermis and reduced density of dermal collagen. Whilst UV-exposed skin from the same donor identified an increased number of melanocytes which were more pronounced in older donors, and coincided with dermal collagen structure disturbances. Differential expression in mixed sex samples identified over-representation of TGF-beta SMAD regulated genes (ATOH8, ID3, ID1, SMAD7, FAM83G) which were highly correlated. Following this sex specific expression profiles were assessed, which identified 57 per cent of the genes had highest expression in males and 43 per cent had highest expression in females. Functional enrichment of these genes revealed over-representation of metabolic steroid biosynthesis^9^. Table 1 provides an overview of studies that have previously analysed fibroblast and skin data assessed in this study, which builds on previous findings by stratifying samples by previously identified factors (age, sex, UV-exposure status)^9, 12^.

**Table 1.**
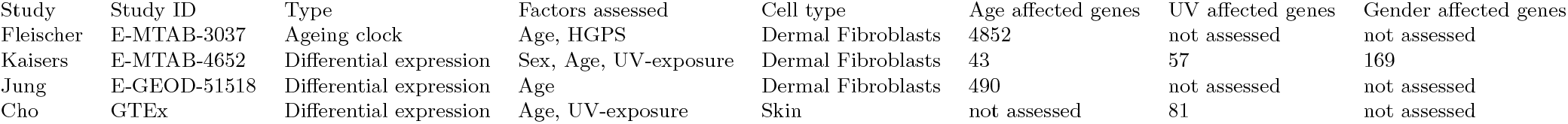
Overview of ageing studies previously completed using the data sets assessed in this study, factors assessed and the number of differentially expressed genes identified

The GTEx skin RNA-seq data analysed in this study has previously been analysed in the context of UV exposure by Cho et al. 2018^12^, in total 606 samples were assessed (228 UV-protected and 349 UV-exposed). A mixed sex analysis comparing UV-exposed and UV-protected skin. The results identified decreased expression of histone genes (HIST) and increased expression of MMPs (MMP3, MMP13, MMP2, MMP8, MMP10) in UV-exposed skin. Pathway analysis on genes increased in expression identified enrichment of the epidermal differentiation complex, which regulates skin barrier function and comprises SPRRs, LCEs, S100 genes. Whilst genes that were decreased in expression regulated responses to external stimulus, cell proliferation and lipid metabolism. Lipid metabolic genes (ACSBG1, ALOX15B, ELOVL3, FADS1, FADS2, THRSP) were associated with patients with psoriatic dermatitis, whilst CYP4F8, ELOVL3, FADS1, FADS2, FAR2, HAO2 were decreased in patients with atopic dermatitis. In addition genes increased in expression in psoriatic skin (LCE3D, SPRR2B and SPRR2G) were increased in expression in UV-exposed skin^12^.

### The role of fibroblasts in skin maintenance

Fibroblasts located in the extracellular matrix (ECM) produce a sophisticated network of structural substances (collagen and elastin), proteoglycans, mucopolysaccharides and cell adhesion proteins (Glycosaminoglycans (GAGs) hyaluronic acid and chondroitin, that maintain hydration), providing structural and biochemical support to the ECM and surrounding cells^1, 3, 7–9^. Fibroblasts have been widely studied in attempts to understand skin ageing; Kaisers did not detect age-related alterations in the transcriptomes of short term cultured fibroblasts^9^, whilst Fleischer successfully used machine learning to predict the age of donors from transcriptomic data^11^. In reduced lipid conditions miRNAs identified altered actin dynamics, proinflammatory cytokines, and TGF-beta signal transduction^13^. Aged fibroblasts undergo a reduction in metabolic activity, resulting in reduced synthesis, degraded and abnormally located fibres, the accumulation of senescent fibroblasts with reduced apoptosis, that do not orient along elastin fibres^1, 4, 7–9^. These changes are thought to be linked to altered expression of matrix metalloproteinases (MMPs)^4^; linear declines of 29 per cent over 49 years in collagen I have been associated with elevated MMP1^9^. A diagrammatic overview of the ECM and ECM-fibroblast interactions is provided in Figure 1.

**Figure 1.**
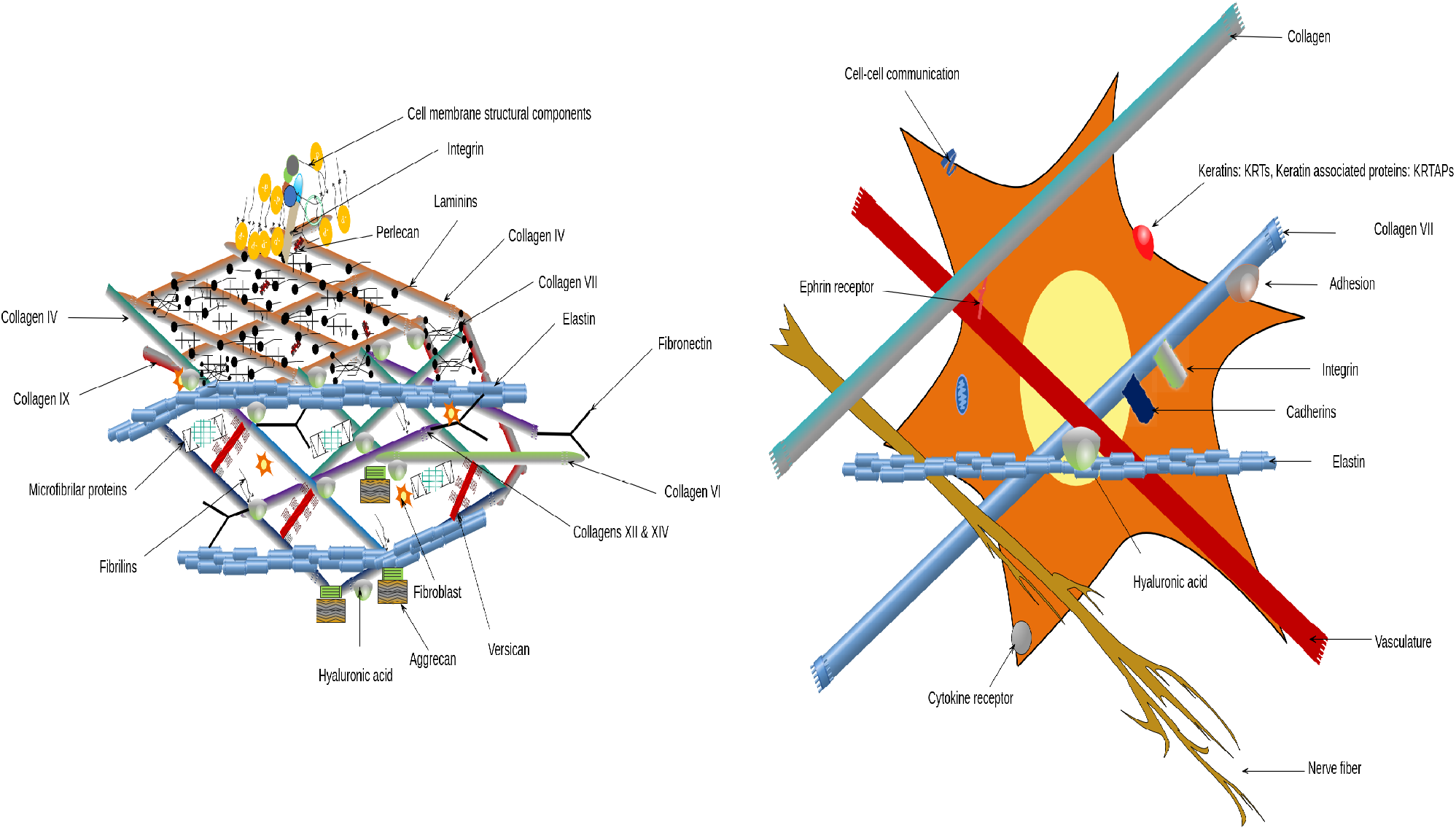
Molecular diagram of a young healthy ECM and fibroblasts identifying key structures, genes, proteins, processes and interactions involved in maintaining skin structure. Generated using references^1–8, 14–21^

**Figure 2.**
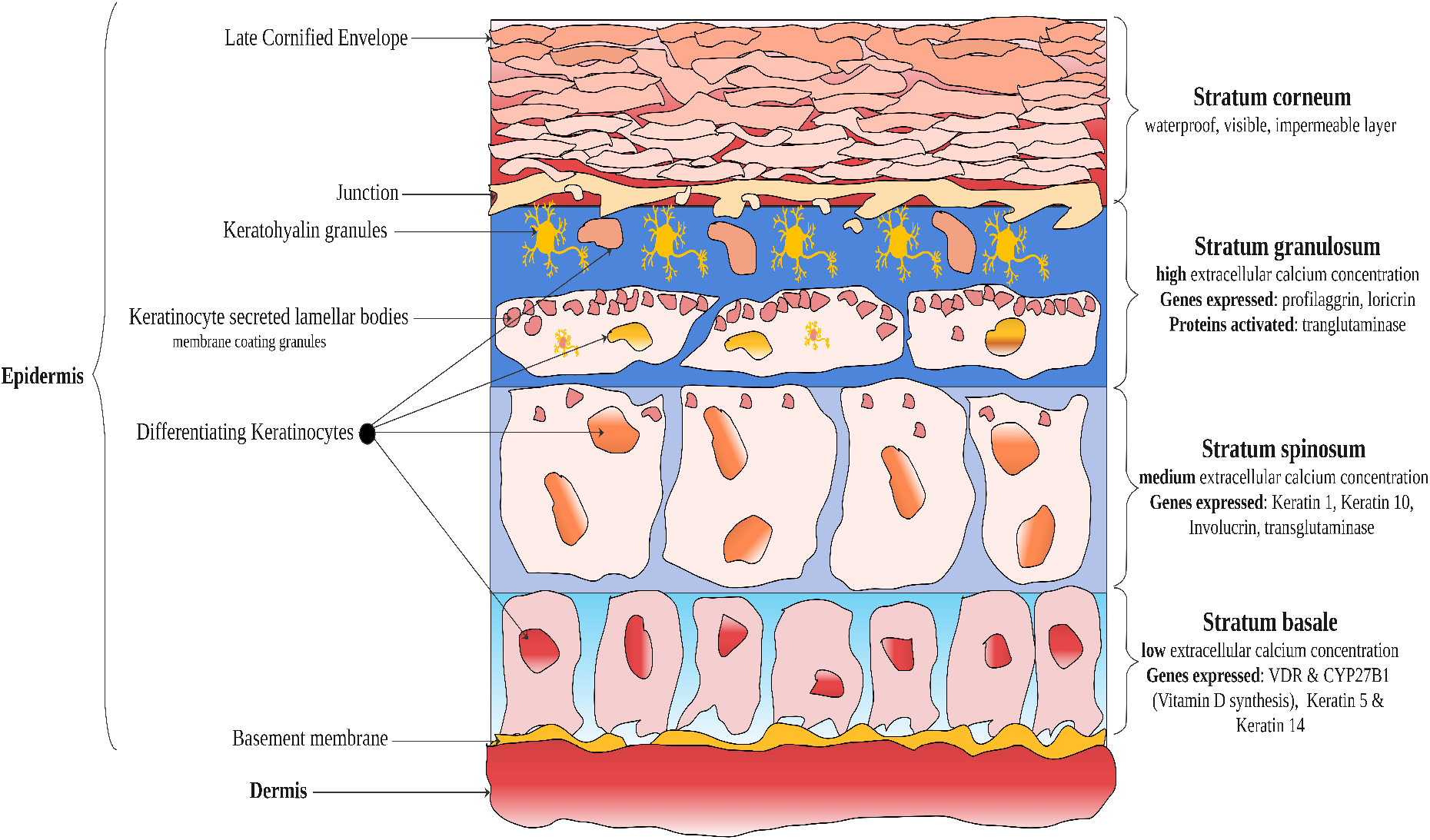
Diagram of the epidermis showing the four layers and key features (calcium concentrations, genes expressed, and proteins activated) which facilitate the differentiation of keratinocytes to form healthy skin. Generated using references^1–4, 7, 9, 14–21^

**Figure 3.**
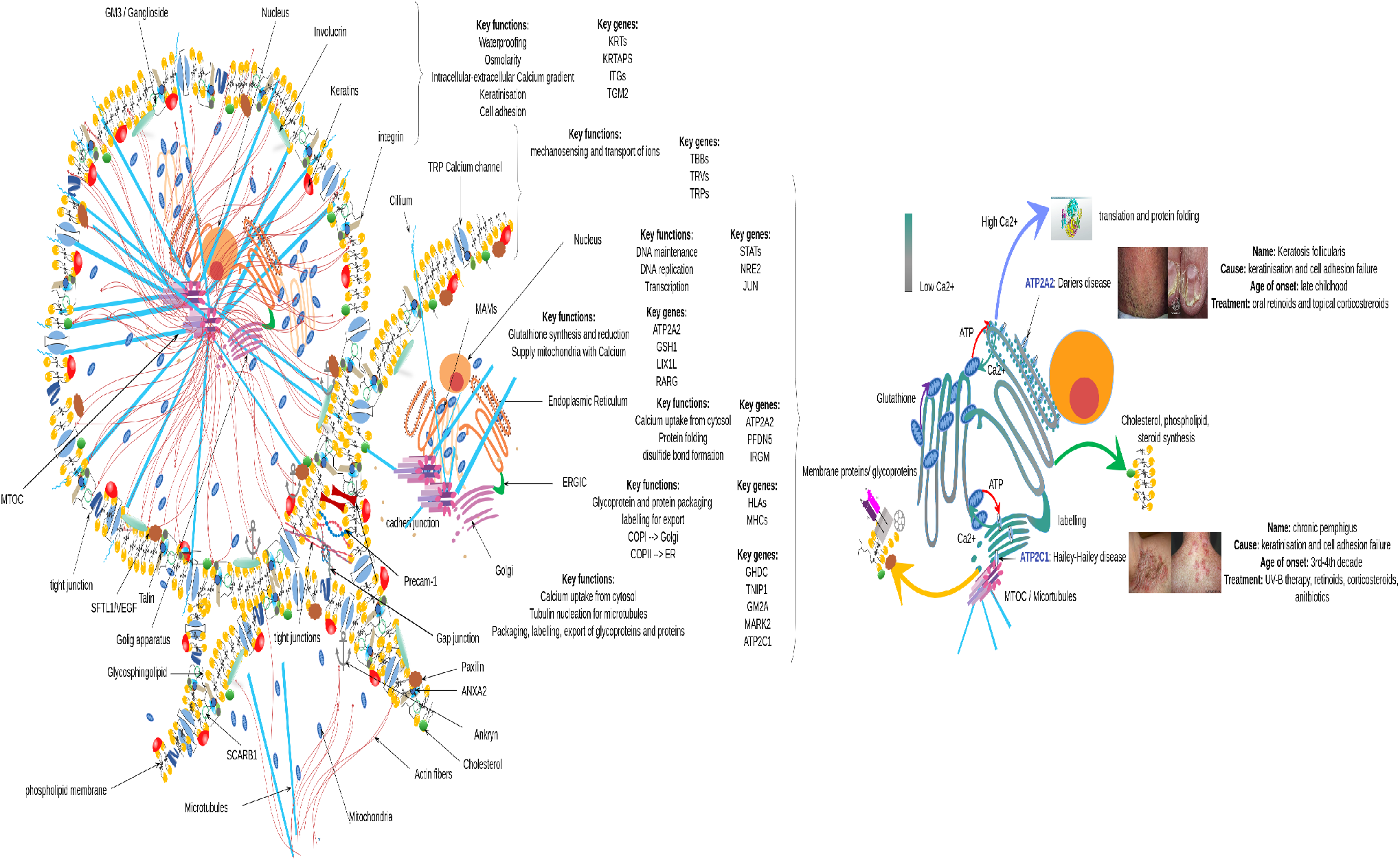
Molecular diagram of a young healthy keratinocyte highlighting key organelles (MAMs, ER, golgi apparatus), genes and processes (left) and consequences of loss of function mutations in ATP2A2 and ATP2C1 in skin development (right). Generated using references^1–4, 7, 9, 14–29^

**Figure 4.**
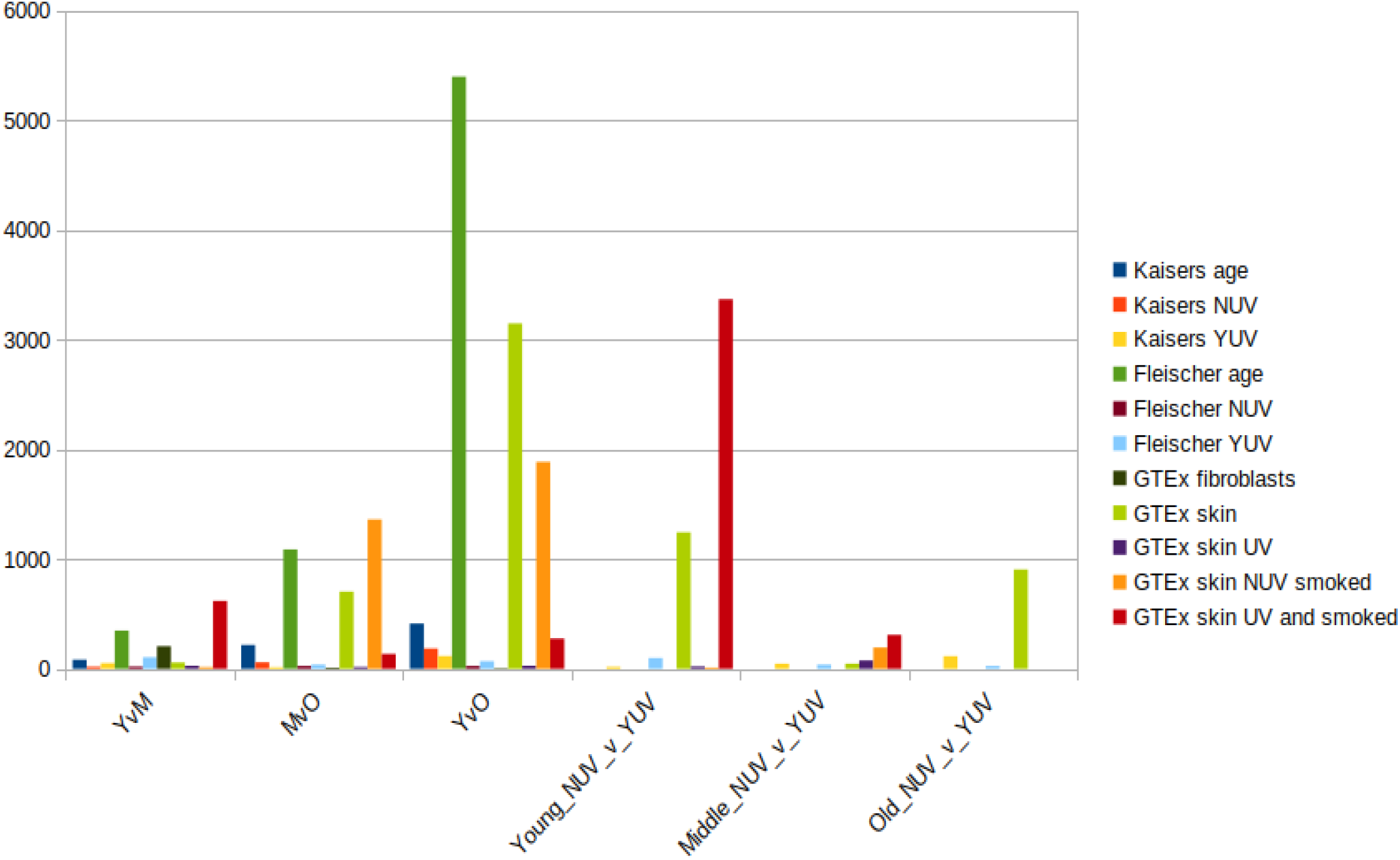
Barchart of the number of significantly (q< 0.05) DE genes identified using Salmon for male age group comparisons of fibroblasts (Kaisers, Fleischer, GTEx fibroblasts) and skin (GTEx skin) with and without environmental stratification’s

Transcriptomic assessment of fibroblasts isolated from patients and cultured for two passages has identified increases in the expression of genes involved in cell adhesion, collagen and extracellular matrix organisation alongside visible alterations in cell and micro-fibril organisation. Subsequent transplantation of cultured fibroblasts improved dermal morphology and skin had more numerous fibroblasts, thicker collagen fiber bundles, higher density of micro-fibrils and elastin fibers after three months^4^. Fibroblast Growth Factor (FGF) has been shown to activate the synthesis of elastin and collagen forming the extracellular matrix leading to increased skin smoothness, thickness and viscoelasticity with pre and post treatment age differences of more than 20 years^7^. Thus far skin ageing treatments have focused on application of topical hyaluronic acid, collagen, as well as injection of soft tissue fillers, hyaluronic acid, collagen and autologous fat, and dietary supplementation with the glycosaminoglycan chondroitin sulphate (involved in hydration, growth factor and cytokine binding), which target and promote rejuvination of the ECM^3, 4^. It is therefore possible that the ageing process in skin is driven by the fibroblast micro-environment; culturing the cells corrects for this. Complications in assessing ageing signatures in cultured fibroblasts can therefore arise because the passage at which samples are assessed can influence methylation and transcriptional profiles^13^, as well as physical processes such as the uptake of Ca^2+^ by mitochondria which can induce senescence; these effects can also differ from one cell type to another^22^.

### Epidermal structure and maintenance

The epidermis comprises four functionally different layers (stratums; basale, spinosum, granulosum, corneum) of keratinocytes in different stages of differentiation^2^. An overview of epidermal structure and conditions regulating the differentiation of keratinocytes is provided in 2.

Briefly the stratum basale; the innermost layer, has low extracellular Ca^2+^ concentrations which regulate synthesis of vitamin D in response to UV-exposure, produce keratin 5 and 14^2, 16^. In the stratum spinosum, extracellular Ca^2+^ increases triggering the expression of keratins 1, 10, transglutaminase and involucrin^2, 16^. The stratum granulosum has the highest Ca^2+^, here proflaggrin, loricrin are produced in keratohyalin granules, lipids are added for waterproofing, whilst transglutaminase is activated, cross-linking involucrin to the cell membrane^2, 16^. The stratum corneum is comprised of the cornified envelope, attached to the lipid envelope secreted by the lamellar bodies in the stratum granulosum^2, 16^ resulting in a resilient impermeable structure protecting the viable layers underneath^2, 16^. Mutations in, or absence of structural proteins such as keratins (5/14, 1/10, 6/16,17,9), filaggrin, the absence or inactivation of Ca^2+^ dependent cross linking enzyme (transglutaminase) are implicated in a range of skin disorders^17^. The calcium gradient between the stratum basale and the stratum spinosum regulates differentiation of keratinocytes into skin layers; achieved by calcium and the active metabolite of vitamin-D which sequentially turn on and off the genes required to produce the elements for differentiation^16^. When the calcium gradient is lost, lamellar body secretion increases, and expression of differentiation markers (loricrin, proflaggrin and involucrin) decrease^2^, leading to a reduction in cross-linking and structural integrity of the stratum corneum^17^, restored by exposure to extracellular calcium^2^. In normal keratinocytes increased intracellular calcium stimulates the formation of cell-cell adherens junctions, and reorganisation of the actin cytoskeleton^18^ as well as regulating the distribution and solubility of cadherins and integrins^21^. It has thus been established that calcium gradients in conjunction with vitamin-D regulate the differentiation status of keratinocytes, and that disruption of calcium ATP pumps (ER, Golgi) lead to skin defects^2, 15–17^. Cultured human epidermal keratinocytes increase cadherins, integrin, alpha-catenin, beta-catenin, plakoglobin, viniculin and alpha-actinin in response to increases in extracellular Ca^2+^^21^. The concentrations of Ca^2+^ are tightly controlled by translocation of PKC to cell membranes, opening of calcium channels, increasing intracellular Ca^2+^; transglutaminase is activated at high Ca^2+^ in the stratum granulosum^16^. Cytosolic Ca^2+^ is precisely regulated because it controls essential cell functions such as proliferation, differentiation, secretion, contraction, metabolism, trafficking, gene transcription and apoptosis^15, 23–25^.

### The role of Calcium gradients in keratinocyte and skin development

Disruption of calcium ATP pumps (ER, Golgi) lead to skin defects^2, 15–18^; mutations in the Golgi calcium pump (ATP2C1) leads to altered intracellular calcium, due to disrupted transport into the golgi apparatus leading to Hailey-Hailey disease^18^. In Hailey-Hailey disease decreased ATP is observed in conjunction with impairment of the calcium induced reorganisation of actin filaments^18^, which is thought to result from calcium overload in the mitochondria^18^. Onset of Hailey-Hailey disease occurs in middle age, consistent with increased mitochondrial sensitivity to calcium overload^18^. Disruption of the ER calcium pump (ATP2A2/SERCA2) disrupts K1/K10 transitions leads to Darier’s disease, also observed in epidermolytic itchyosis characterised by increased thickness of stratum granulosum, with increased cell divisions^17^. Both inherited conditions are associated with impaired calcium gradients resulting in decreased cell adhesion, decreased Keratin 14 expression and increased expression of Keratin 10^2, 17^. In normal keratinocytes increased intracellular calcium stimulates the formation of cell-cell adherens junctions, and reorganisation of the actin cytoskeleton^18^ as well as regulating the distribution and solubility of cadherins and integrins^21^.

The ER is a an intracellular Ca^2+^ store that regulates cytosolic Ca^2+^ as well as assembling and folding newly synthesised proteins, disruption of ER homeostasis and Ca^2+^ depletion are involved in the pathophysiology of many diseases^15^. To function as an intracellular store of Ca^2+^ the ER expresses three different types of proteins. Sarco/endoplsmic reticulum Ca^2+^ ATPase (SERCA) actively pumps Ca^2+^ into the ER lumen, internal Ca^2+^ is sequestered by calsequestrin, calrecticulin and calnexin, immunoglobulin heavy chain binding protein (BiP) and protein disulfide isomerases (PDI), regulating binding, calcium import and release^15, 23^. These proteins also bind misfolded proteins via inappropriately exposed hydrophobic or hypo-glucosylated residues, PDI mediates the correct formation of disulfide binds, and require high Ca^2+^ for their activity, depletion of ER Ca^2+^ leads to the inappropriate secretion, aggregation and degradation of unassembled proteins^15, 23^.

Mitochondrial associated membranes (MAM) are connected to the ER, they are involved in processes such as lipid biosynthesis and trafficking, calcium homeostasis, reactive oxygen species (ROS) production and autophagy^22^. The membranes provide a physical platform which enables communication between the ER and mitochondria which exerts control over lipid synthesis and Ca^2+^ and lipid transport within and between cells^22^. Mitochondria ER contacts (MERCs) regulate the mitochondrial network and mark the sites of mitochondrial fission, and thus dysregulation of the ER can impact on mitochondrial outputs (ATP) and replenishment^22^.

Uptake of calcium affects the krebs cycle, enzyme activity, ATP synthesis, mitochondrial permeability transition pore opening, and thus mitochondrial membrane potential, respiration and ROS production^15, 22^. Mitochondrial Ca^2+^ overload induces cell death by increasing outer membrane permeability resulting in loss of the mitochondrial membrane potential^15^. Age related declines in function of MAM have been associated with age related degenerative disorders including Alzheimers, lateral sclerosis, type 2 diabetes, obesity, GM1 gangliosidosis and viral infections including cytomegalovirus and hepatitis C^22^. Ageing is accompanied by increased oxidative damage and the dysfunction of ER proteins^22^, MAM are thought to play a role in the regulation of ROS production by ER and mitochondria^22^; the rate of production of ROS depends on the redox state of the respiratory chain, the status of MAM, all of which depend on Ca^2+^ homeostasis^22^. An diagrammatic overview of the molecular structure of a cell, the visible consequences of loss of function mutations in ATP2A2 and ATP2C1 on the MAM-ER-Golgi-ERGIC-cytoskeletal complex are shown in 3

Calcium is well known for it’s structural role in the formation of bones and teeth, however calcium is also an important structural component of individual cells and the extracellular matrix. It acts as a cofactor required for the stabilisation of disulfide bonds between peptides for structural proteins including; collagen^20^, cell-cell adhesion molecules such as integrin^30^, and cell-cell junctions formed by cadherins^31^. In addition it plays a vital role in formation of the cytoskeleton; microtubule elongation from nucleation of tubulin^25^, actin-fiber from globular actin^24, 32^. Once inside the cell calcium accumulates in golgi apparatus and ER against a calcium gradient; mediated by APT2A2 (ER) and ATP2C1 (golgi); hydrolysing ATP supplied by the mitochondria and in turn supplying Ca^2+^ to the mitochondria^15, 23^. Concentrating different amounts of Ca^2+^ in the ER and golgi apparatus generates compartments with differing pH (ER=(7.1-7.4), golgi=(5.9-6.3), mixing of their fluids, and control over flow through the ER Golgi Intermediate Compartment (ERGIC) allows for precise control over pH, calcium availability and thus protein folding and extension conditions^15, 20, 33^.

Proteins sensitive to Ca^2+^ transduce key signalling pathways which allow cells to respond appropriately to a range of stimuli such as depolarisation of cell membranes, osmolarity, cell surface distortion and changes in temperature^24^. Specific channels sensitive to hormones and cytokines line cell membranes and are highly coordinated controlling the amplitude and duration of cytosolic Ca^2+^ spikes^23–25^. Calcium induces the src family tyrosine kinases, activation of PLC-y1^16^, whilst calcium stress inhibits mRNA translation to reduce the influx of new proteins into the ER, and activation of MAPK signalling^15^. Release of Ca^2+^ from the ER, combined with influx from extracellular space leads to increased cytosolic calcium levels and the activation of Protein Kinase C (PKC), calcineurin, calpains, calmodulin dependent kinases and calmodulin binding proteins^23^. In this respect calcium is able to regulate the cell cycle, consequently calcium dysregulation has been observed in some cancers (colon carcinoma, Myeloid leukemia, Breast cancer)^23^.

Skin maintenance and ageing are affected by a wide range of factors; targeting therapies at molecules and proteins known to be reduced, regulatory hormones/growth factors, extracting, culturing and re-introducing fibroblasts have all proven successful. Analysis complications can arise because the passage at which samples are assessed can influence methylation and transcriptional profiles^13^, as well as physical processes such as the uptake of Ca^2+^ by mitochondria which can induce senescence; these effects can also differ from one cell type to another^22^.

Here we aimed to determine the impact of ageing on whole skin and fibroblasts specifically in males at the transcriptional and epigenetic level. Alignment free quantification (Salmon) of publicly available mRNA from cultured fibroblasts determined factors, (biological, clinical and experimental) influencing gene expression. The research was extended to include skin and uncultured fibroblasts from the extensively annotated GTEx repository, and the impacts of UV-exposure and smoking status were assessed using two tools; Salmon and CuffDiff. The research further aimed to identify related epigenetic alterations in ageing male skin tissues.

## Results

To identify genes associated with ageing in human fibroblasts we used a semi-systematic approach (see methods) to include studies with RNA-seq data for fibroblasts from individuals at different ages. Additional inclusion and exclusion criteria are described in the methods. We identified three studies with publicly available cultured fibroblast RNA-seq data which was analysed in conjunction with data from the GTEx repository uncultured fibroblasts. The GTEx repository also contained RNA-seq data for skin, which offered an opportunity to compare and contrast age related transcriptomic changes in skin with those seen in cultured fibroblasts, Table 2.

**Table 2.**
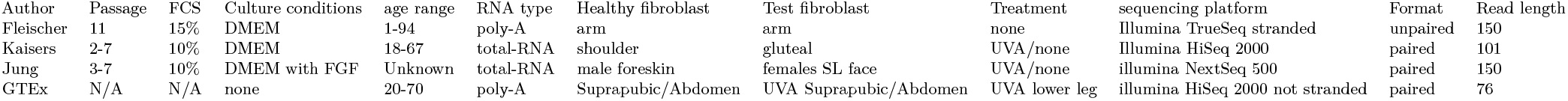
Summary of key differences in experimental conditions for each of the RNA-seq studies conducted on aged fibroblasts. Abbreviations FBS=Fetal Bovine Serum, FGF= Fibroblast Growth Factor

| Author | Passage | FCS | Culture conditions | age range | RNA type | Healthy fibroblast | Test fibroblast | Treatment | sequencing platform | Format | Read length |
| --- | --- | --- | --- | --- | --- | --- | --- | --- | --- | --- | --- |
| Fleischer | 11 | 15% | DMEM | 1-94 | poly-A | arm | arm | none | Illumina TrueSeq stranded | unpaired | 150 |
| Kaisers | 2-7 | 10% | DMEM | 18-67 | total-RNA | shoulder | gluteal | UVA/none | Illumina HiSeq 2000 | paired | 101 |
| Jung | 3-7 | 10% | DMEM with FGF | Unknown | total-RNA | male foreskin | females SL face | UVA/none | illumina NextSeq 500 | paired | 150 |
| GTEx | N/A | N/A | none | 20-70 | poly-A | Suprapubic/Abdomen | UVA Suprapubic/Abdomen | UVA lower leg | illumina HiSeq 2000 not stranded | paired | 76 |

### RNA-seq fibroblast sample attributes, treatments and platforms

RNA-seq data from cultured fibroblasts and skin along with the methods used in data generation are shown in Table 1.

Table 2 identified that there are key differences in the way that cells were cultured prior to RNA extraction. The quantity of Fetal Bovine Serum used in culture medias varied between studies, additionally Jung et al (2018) stimulated the growth of cells using fibroblast growth factor, known to regulate TGF-Beta. These key study differences were used to derive analysis decisions further discussed in the methods.

### Functional Enrichment: Fibroblasts

Since expression profiles revealed fundamental differences in the transcriptomes of fibroblasts and skin, they were analysed separately. In addition each of the data sets was analysed independently because they represented different age ranges, male female ratios and culturing conditions differed. An overview of the number of differentially expressed genes and functional enrichment categories identified in male age group comparisons of fibroblasts in each of the data sets is presented in Table 3.

**Table 3.** Overview of the number of genes and enrichment categories identified for GTEx uncultured male fibroblast age group comparisons and male specific re-analysis of Fleischer and Kaisers data sets by age group assessed using Salmon

| Comparison | Genes | Pathways | GOBP | GOCC | GOMF |
| --- | --- | --- | --- | --- | --- |
| GTEx_Young_vs_Middle_males | 207 | 0 | 0 | 0 | 0 |
| Fleischer_Young_vs_Middle_males | 348 | 0 | 0 | 0 | 0 |
| Kaisers_Young_vs_Middle_males | 84 | 0 | 0 | 0 | 0 |
| GTEx_Middle_vs_Old_males | 12 | 0 | 0 | 0 | 0 |
| Fleischer_Middle_vs_Old_males | 1088 | 6 | 0 | 4 | 2 |
| Kaisers_Middle_vs_Old_males | 217 | 0 | 0 | 0 | 0 |
| GTEx_Young_vs_Old_males | 6 | 0 | 0 | 0 | 0 |
| Fleischer_Young_vs_Old_males | 5397 | 15 | 75 | 33 | 14 |
| Kaisers_Young_vs_Old_males | 413 | 0 | 0 | 0 | 0 |

Table 3 identifies that most genes are identified in the Fleischer data set age group comparisons, and this corresponds to increased enrichment. A summary of the top enrichment categories by significance is shown in Table 4.

**Table 4.** Overview of the top five enrichment categories by significance identified from Fleischer cultured male fibroblast young vs old age group comparisons of males assessed using Salmon

| Category | ID | Description | GeneRatio | BgRatio | qvalue | Comparison |
| --- | --- | --- | --- | --- | --- | --- |
| BP | GO:0043269 | regulation of ion transport | 19/162 | 129/3679 | 0.004464542915939 | YvO |
| BP | GO:0007186 | G protein-coupled receptor signaling pathway | 20/162 | 149/3679 | 0.005223144902447 | YvO |
| BP | GO:1901342 | regulation of vasculature development | 13/162 | 75/3679 | 0.006333206761025 | YvO |
| BP | GO:2000026 | regulation of multicellular organismal development | 41/162 | 488/3679 | 0.006333206761025 | YvO |
| BP | GO:0001525 | angiogenesis | 17/162 | 123/3679 | 0.006333206761025 | YvO |
| MF | GO:0030545 | receptor regulator activity | 18/158 | 97/3645 | 2.28425240129393E-05 | YvO |
| MF | GO:0005102 | signaling receptor binding | 35/158 | 342/3645 | 7.53215642493093E-05 | YvO |
| MF | GO:0008201 | heparin binding | 11/158 | 42/3645 | 7.53215642493093E-05 | YvO |
| MF | GO:0004930 | G protein-coupled receptor activity | 13/158 | 64/3645 | 0.00013209571385 | YvO |
| MF | GO:0004888 | transmembrane signaling receptor activity | 18/158 | 131/3645 | 0.000447215841956 | YvO |
| CC | GO:0031012 | extracellular matrix | 18/167 | 122/3831 | 0.001021000902685 | YvO |
| CC | GO:0005887 | integral component of plasma membrane | 28/167 | 271/3831 | 0.001454977112649 | YvO |
| CC | GO:0031226 | intrinsic component of plasma membrane | 28/167 | 287/3831 | 0.00287285466026 | YvO |
| CC | GO:0062023 | collagen-containing extracellular matrix | 14/167 | 99/3831 | 0.005257901207925 | YvO |
| CC | GO:0009986 | cell surface | 18/167 | 160/3831 | 0.008713090648965 | YvO |
| KEGG | hsa04080 | Neuroactive ligand-receptor interaction | 5/24 | 11/361 | 0.007593134990236 | YvO |

**Table 5.** Overview of the number of genes and enrichment categories identified for GTEx male skin age group comparisons, assessed using Salmon, and the number of differentially methylated CpGs identified (DM CpGs) and enrichment categories for corresponding age group comparisons

| Comparison | Genes/CpGs | Pathways | GOBP | GOCC | GOMF |
| --- | --- | --- | --- | --- | --- |
| Young_vs_Middle_males | 56 | 1 | 20 | 16 | 4 |
| DM_CpGs_Young_vs_Middle_males | 1 | 0 | 0 | 0 | 0 |
| Middle_vs_Old_males | 703 | 8 | 0 | 0 | 0 |
| DM_CpGs_Middle_vs_Old_males | 110998 | 65 | 300 | 99 | 67 |
| Young_vs_Old_males | 3149 | 6 | 193 | 42 | 52 |
| DM_CpGs_Young_vs_Old_males | 162268 | 66 | 773 | 100 | 104 |

**Table 6.** Overview of the top five enrichment categories by significance identified for GTEx male skin young vs old age group comparisons assessed using Salmon

| Category | ID | Description | GeneRatio | BgRatio | qvalue | Comparison |
| --- | --- | --- | --- | --- | --- | --- |
| BP | GO:0031424 | keratinization | 9/116 | 12/2052 | 1.49702764472407E-06 | YvO |
| BP | GO:0009888 | tissue development | 35/116 | 230/2052 | 7.15878243115328E-06 | YvO |
| BP | GO:0051240 | positive regulation of multicellular organismal process | 32/116 | 212/2052 | 3.20084787748422E-05 | YvO |
| BP | GO:0030154 | cell differentiation | 53/116 | 497/2052 | 7.03795496677523E-05 | YvO |
| BP | GO:0030216 | keratinocyte differentiation | 9/116 | 20/2052 | 0.00015537034513 | YvO |
| MF | GO:0048018 | receptor ligand activity | 12/107 | 45/2040 | 0.000301285992384 | YvO |
| MF | GO:0030545 | receptor regulator activity | 12/107 | 48/2040 | 0.000319370615109 | YvO |
| MF | GO:0005125 | cytokine activity | 8/107 | 21/2040 | 0.000324811953653 | YvO |
| MF | GO:0005102 | signaling receptor binding | 24/107 | 188/2040 | 0.00089492601771 | YvO |
| MF | GO:0005201 | extracellular matrix structural constituent | 10/107 | 42/2040 | 0.001420235367856 | YvO |
| CC | GO:0031012 | extracellular matrix | 21/116 | 96/2150 | 2.04081127836822E-06 | YvO |
| CC | GO:0005882 | intermediate filament | 8/116 | 16/2150 | 4.75374246898633E-05 | YvO |
| CC | GO:0072562 | blood microparticle | 10/116 | 28/2150 | 4.75374246898633E-05 | YvO |
| CC | GO:0044421 | extracellular region part | 47/116 | 457/2150 | 4.75374246898633E-05 | YvO |
| CC | GO:0045111 | intermediate filament cytoskeleton | 8/116 | 20/2150 | 0.000142007739314 | YvO |
| KEGG | hsa04080 | Neuroactive ligand-receptor interaction | 10/57 | 33/973 | 0.000601826946539 | YvO |
| KEGG | hsa04024 | cAMP signaling pathway | 7/57 | 16/973 | 0.000601826946539 | YvO |
| KEGG | hsa04060 | Cytokine-cytokine receptor interaction | 8/57 | 30/973 | 0.005819688427756 | YvO |
| KEGG | hsa04610 | Complement and coagulation cascades | 6/57 | 20/973 | 0.015364147000617 | YvO |

**Table 7.**
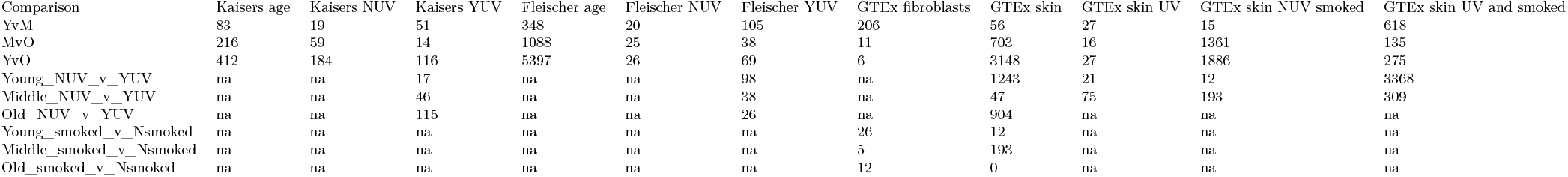
Overview of the number of DE genes identified using Salmon for all possible comparisons (na means a comparison was not possible due to lack of annotation or insufficient samples) of fibroblasts and skin samples

Table 4 identifies that in fiborblasts the top GO biological process is regulation of ion transport, whilst molecular functions represent signalling and receptor activities, cellular component categories relate to the extracellular matrix and plasma membrane, which leads to the identification of the neuroactive ligand-receptor interaction pathway.

### Differential expression and methylation in skin

Significantly (q < 0.05) DE genes and transcripts were functionally enriched by identifying enriched Gene Ontology Biological Process categories and KEGG pathways. DM analysis also identified significant age related differences in males that were functionally enriched (Table 8).

**Table 8.** Overview of the levels of stratification leading to the identification of functional enrichment for tools (Salmon = Sa, CuffDiff reference = CDref, CuffDiff *de novo* = CD_dn), for quantification of genes (gns) and isoforms (iso), and Limma for identification of differential methylation in skin

| Comparison | No stratification | UV-exposed | Smoke | Non UV-exposed smoked |
| --- | --- | --- | --- | --- |
| Young v Middle male | CDref_iso, CDref_gns, CD_dn_iso | CD_dn_iso, CD_dn_gns, Sa | Sa | Sa |
| Middle v Old male | Sa, Limma, CD_dn_iso, CD_dn_gns | CD_dn_iso | 0 | Sa |
| Young v Old male | Sa, Limma | CD_dn_gns | 0 | Sa |

Table 5 identifies that most genes and DM CpGs are differentially expressed in young vs old age group comparisons, this age group comparison also leads to the most functional enrichment. An overview of the top functional enrichment categories for youg vs old age group comparisons is shown in 6

The overview of functional enrichment categories in Table 6 identifies that in skin, the most significant biological processes relate to keratinisation and skin development, whilst molecular functions identify signalling and receptor activity, specifically those relating to cytokines. Top cellular component categories relate to the extracellular matrix, blood microparticles, intermediate filaments and the cytoskeleton, whilst the top KEGG category is the same as in fibroblasts; Neuroactive ligand-receptor interaction. Other KEGG categories are reflective of these changes, identifying cAMP signalling, cytokine signalling (involved in viral responses), and blood formation and maintenance (Complement and coagulation cascade).

### Environmental impacts

UV-exposure and smoking are factors known to accelerate skin ageing^1, 3–7, 12, 34–37^, so GTEx data for skin samples was stratified by age group, then UV-exposure and smoking status. Adding additional environmental data improved the number of results obtained for both fibroblasts and skin, identifying that as many genes are differentially expressed in within age group comparisons of UV and non UV-exposed fibroblasts as in age group comparisons of fibroblasts from Fleischer and Kaisers studies.

Independent analysis of each of the fibroblast data sets identified DE genes, with the most genes identified as DE in Fleischer young vs old age group comparisons of fibroblasts (4). When UV exposure status was considered using within age group comparisons, and smoking kept constant (where known for GTEx data) over 3,000 genes were identified as DE in young UV-exposed skin from smokers compared to non UV-exposed skin from smokers 4.

These results highlight the complexity of the underlying data, and the importance of considering known environmental variables during analysis.

### Functional Enrichment in skin: Tools and stratification

An overview of the stratification, datasets and tools which identified functional enrichment categories for skin is summarised in Table 8

Genes and isoforms identified as DE for each age group comparison using CuffDiff and Salmon (CD and Sa) are shown in Table 8. Enriched methylation changes (Limma) are only seen in middle vs old and young vs old age group comparisons.

### Functional enrichment in skin ageing; overview

When skin was assessed independently from fibroblasts, DE genes and transcripts were identified as significantly (q< 0.05) DE for all age group comparisons using both CuffDiff and Salmon (Table 8). An overview of the number of age affected genes and significant functional enrichment categories identified from them is shown in 9. Table 9 identifies that only 59 genes are increased and 103 decreased in ageing male skin, for genes increased in expression in middle and old age immune response categories are enriched.

**Table 9.**
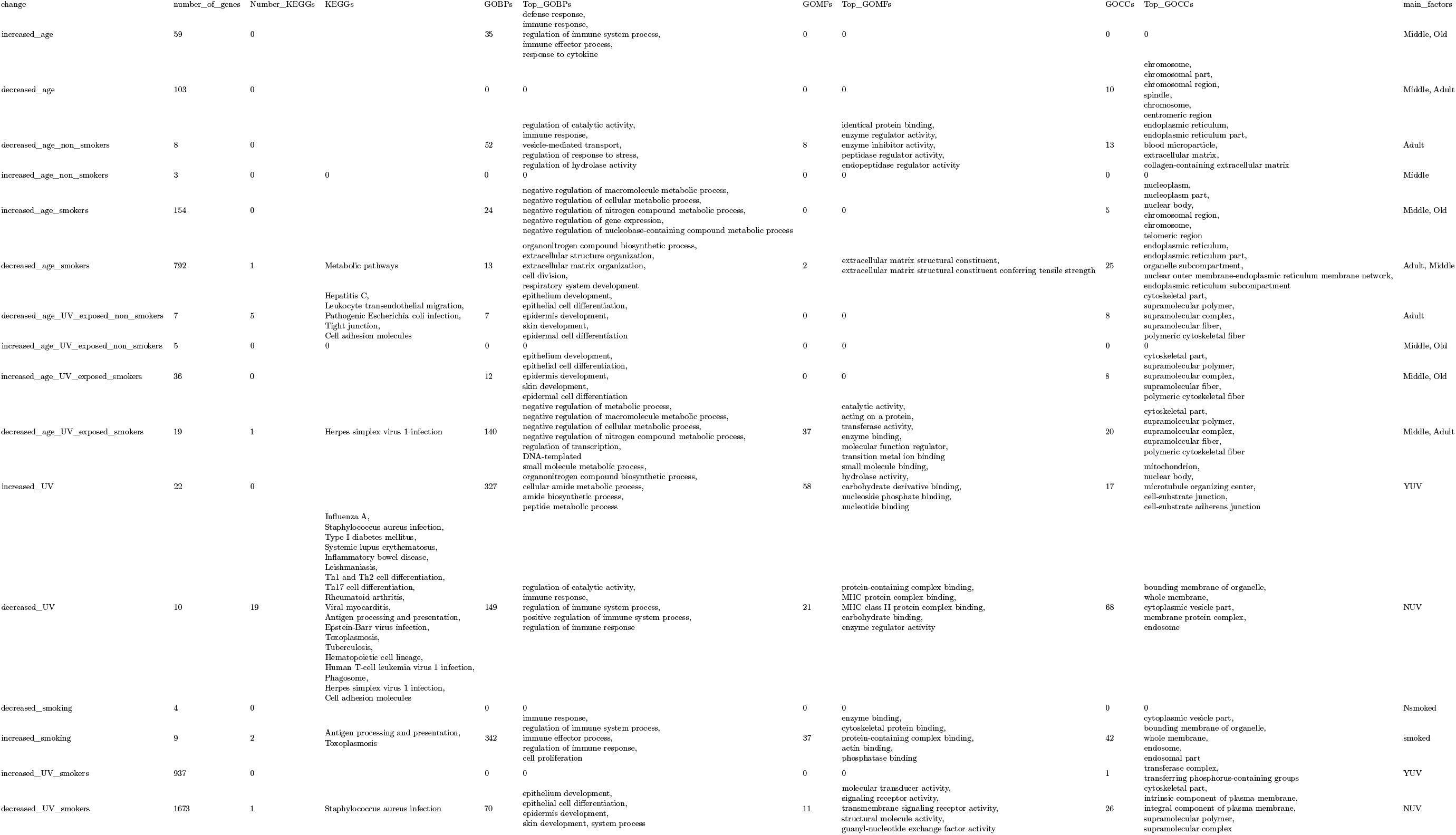
The number of genes and corresponding functionally enriched categories identified for genes DE under combinations of age and environmental conditions assessed, and those increased by UV exposure and smoking

The overview in 9 shows that when samples are stratified by known environmental conditions more genes are identified as DE, and an increase in functional enrichment is observed. In smokers genes with highest expression in young age involved in regulation of catalytic activity, immune responses and regulation of hydrolase activity, and these changes are associated with enzyme regulation occurring the in ER, ECM and blood particles. In UV-exposed skin from non-smokers the highest expression of genes involved in skin development and cytoskeletal development were seen in young adults, interestingly in UV-exposed smokers these genes increased with age, and had higher expression levels in middle and old aged adults. Whilst those decreased in expression in UV-exposed skin from non-smokers were involved in skin development, formation of the cytoskeleton, and supramolecular molecules and complexes, which were increased in expression in middle and old aged UV-exposed smokers. In old aged UV-exposed smokers decreases in negative regulation of metabolism and transcription were seen alongside decreased catalytic activity affecting supramolecular polymers, complexes and cytoskeletal parts. When UV-exposure is considered alone an increase in nitrogen compound synthesis including peptides and amides occurs alongside increased small molecule binding, hydrolase activity, nucleotide binding and these changes are associated with the mitochondria, nucleus, cell junctions and cytoskeleton. UV-exposure decreased the expression of genes involved in response to viral and bacterial infections, regulation of catalytic activity, enzyme regulation, affecting processes in membranes, organelles and the cytoplasm. Smoking increases expression of immune responses, cell proliferation, enzyme, actin, phosphate binding and cytoskeletal protein binding, and these changes are associated with vesicles, organelle membranes and endosomes. Interestingly 9 also identifies some confounding variables, for instance in UV-exposed non smokers skin development categories are decreased with age, whilst in smokers they are increased with age, and when UV-exposed smokers of the same age are compared (decreased UV smokers) UV-exposed skin had lowest expression of skin development genes, thus the data indicates that in smokers UV-exposure decreases the expression of skin development genes. Whilst similar the categories are identified as significant from the DE genes, the genes DE under different conditions were different (Table 10), and thus the impacts may also differ.

**Table 10.**
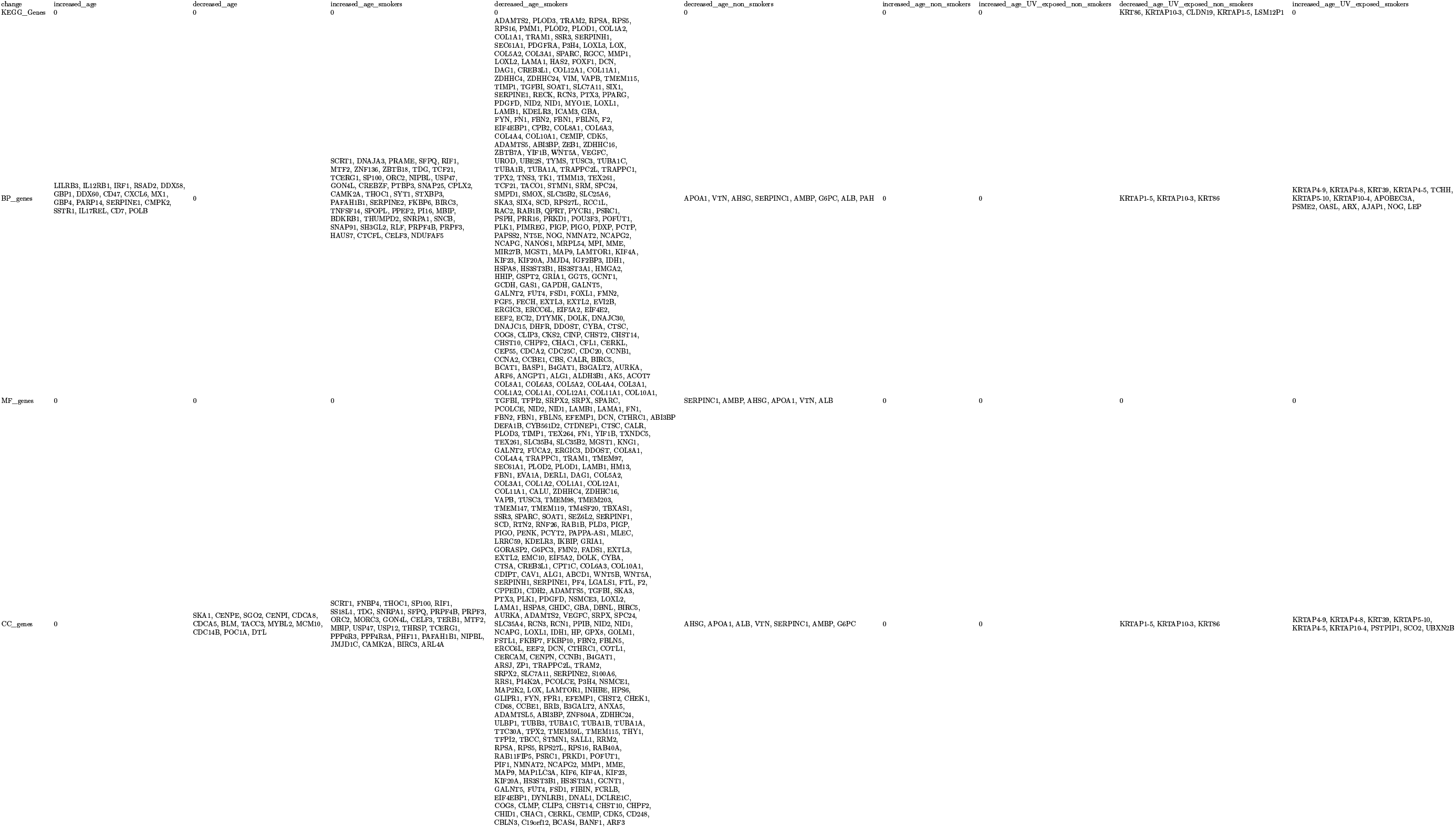
Genes functionally enriched in categories identified for genes DE under combinations of age and environmental conditions assessed, and those increased by UV exposure and smoking

**Table 11.**
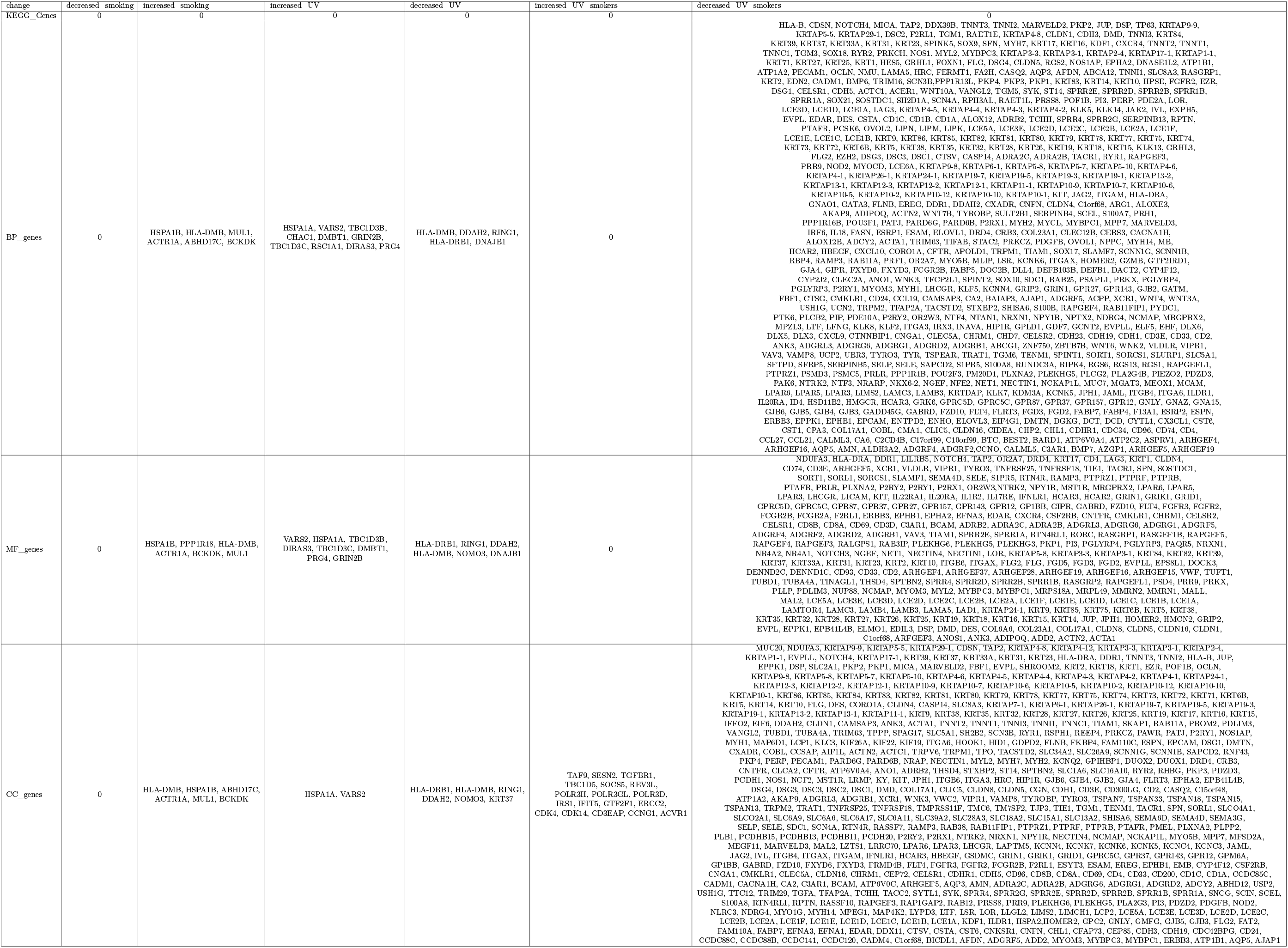
Genes functionally enriched in categories identified for genes DE in response to UV exposure and smoking, and DE in response to UV-exposure and smoking

Table 10 identifies that in the age interpretations some genes increased in expression, whilst genes decreased were enriched for cellular components (most significantly the mitochondria). Most enrichment was identified for gene decreased with age in smokers, affecting both biological processes and cellular components.

When within age group comparisons of environmental variables are assessed a large number of genes are identified as decreased in expression by UV-exposure in smokers 11, irrespective of age group differences, and these are enriched for biological processes, molecular functions and cellular components.

For clarity, key enrichment results for males are presented for age group comparisons in chronological order (young vs middle Young to Middle age changes; 20-40 vs 40-65 years, middle and young vs old Old age changes; 20-65 vs > 65 years). To aid interpret-ability, the results are subset by changes occurring within the cell Male skin; cell and organelle level changes, in the epidermis Male skin: Epidermal changes, in the dermis comprised of ECM, and fibroblast-ECM interactions Male skin: ECM and fibroblasts. Diagrams presented in the discussion provide an overview of DE genes, isoforms, DM gene associated CpGs. Enrichment categories are overlaid on respective diagrams, located according to the location in the cell/tissue that changes are occurring, coloured to indicate increases or decreases of the genes/processes/pathways allowing the consequences of changes to be visualised.

### Young to Middle age changes; 20-40 vs 40-65 years

Reference based CuffDiff identified significant differences and enrichment in young vs middle, only isoforms and genes identified as significantly (q < 0.05) DE by CuffDiff analysis were enriched (q < 0.05) in GO BP categories for young vs middle aged males, this identified that isoforms are involved in viral transcription, protein targeting to membranes, and mRNA catabolic nonsense mediated decay had higher expression levels in middle aged males (Figure 5).

**Figure 5.**
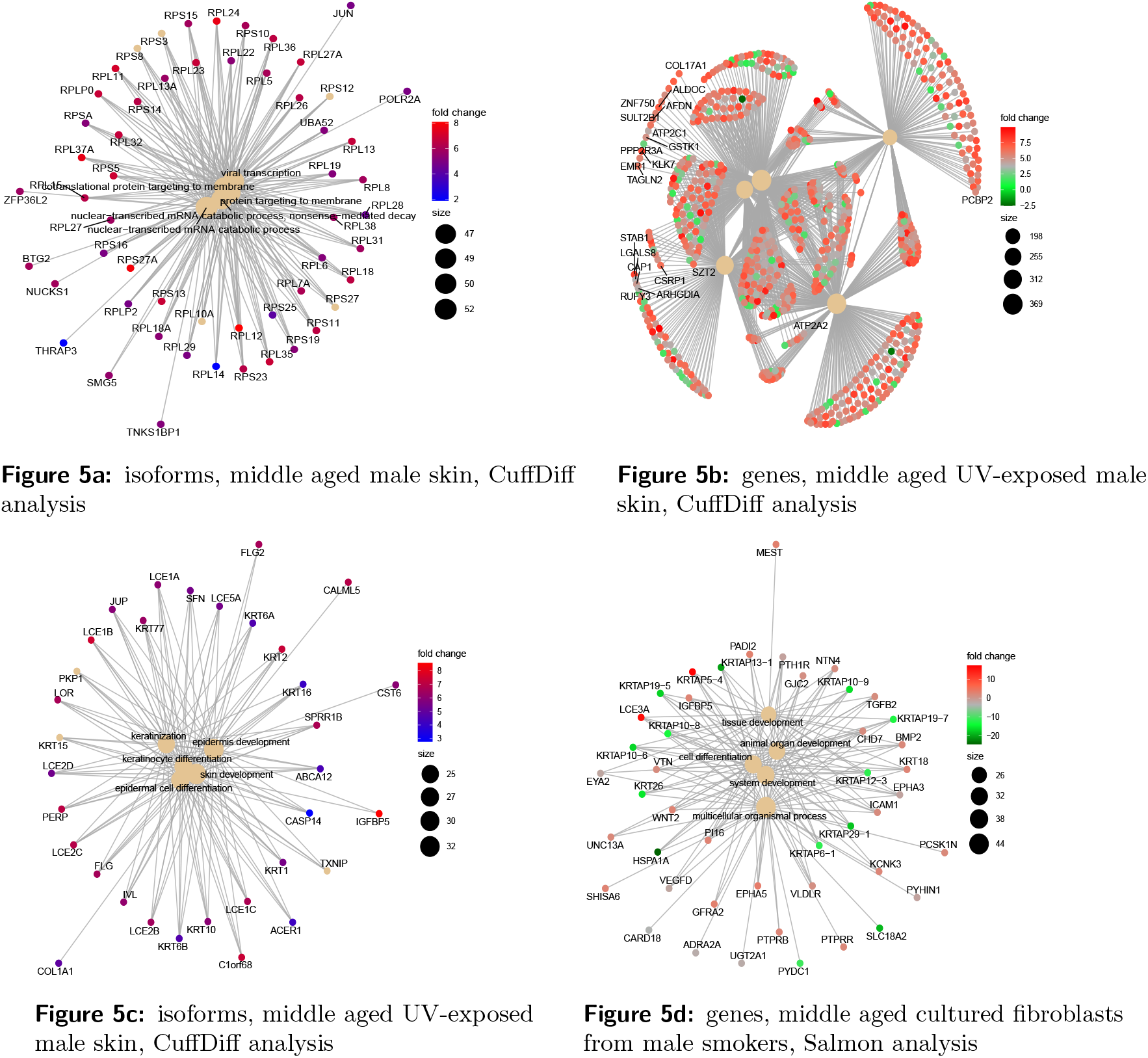
Conserved non-coding element plots represented by significantly (q < 0.05) DE isoforms in middle aged skin (a), significantly (q < 0.05) DE genes (b) and isoforms (c) in young vs middle age group comparisons of UV-exposed male skin; coloured by the log2 fold change in expression in middle aged males. Significantly (q < 0.05) DE genes in male cultured fibroblasts from male smokers (d), coloured by the relative expression level in fibroblasts from young male smokers.

Non-coding elements are conserved sequences within genes which are associated with gene functions, significantly enriched (q < 0.05) conserved non-coding elements for DE isoforms 5Figure 5a identify mostly ribosomal processing (RP) which are consistent with the production of isoforms.

Some additional transcripts of interest include BTG2 (antiproliferative protein regulator of G1/S transition of the cell cycle, JUN, the proto-oncogene, transcription factor and regulator of steroidogenic gene expression). THRAP3 a thyroid hormone receptor associated protein, regulator of lipid metabolism by PPAR alpha. TNKS1BP1 (Tankyrase 1 Binding Protein 1), ankyrin binding, and NUCKS1 (Nuclear Casein Kinase And Cyclin Dependent Kinase Substrate 1). When samples are subsequently stratified by UV-exposure status higher expression of FLG (Filaggrin), LOR (loricin), involucrin (IVL), calmodulin 5 (CALM5), late cornification envelope (LCE), Keratins 1 and 10, (KRT1 and KRT10) isoforms are seen in middle aged UV exposed male skin. Processes relating to skin and epidermal development, keratinisation and keratinocyte differentiation were higher in middle aged male skin exposed to UV 5Figure 5c. Whilst most genes had higher expression in middle aged males, two genes were decreased in expression in middle aged male skin (not shown on plots), these included SPRR2F (associated with keratinisation and Adenosine Phosphoribosyltransferase deficiency), and HLA-DRB5, (a major histocompatibility complex class II, DR beta). Two genes of interest were increased in middle age, ATP2A2 (Darriers disease, an ATP calcium pump in the ER), ATP2C1 (an ATP calcium pump in the golgi, associated with Hailey-Hailey disease) 5Figure 5b, and two genes reflected a change in glycosylation and glutathione production; SULT2B1 a sulfotransferase, GSTK1 a glutathione S-transferase.

### Old age changes; 20-65 vs > 65 years

In old aged males (> 65) significantly DE genes and isoforms were identified using all methods, functionally enriched in GO categories were identified for genes without stratification (Table 8), whilst CuffDiff identified DE genes and isoforms without stratification and in UV-exposed skin.

An overview of functional enrichment for genes and isoforms DE in old aged skin and fibroblasts is shown in Figure 6.

**Figure 6.**
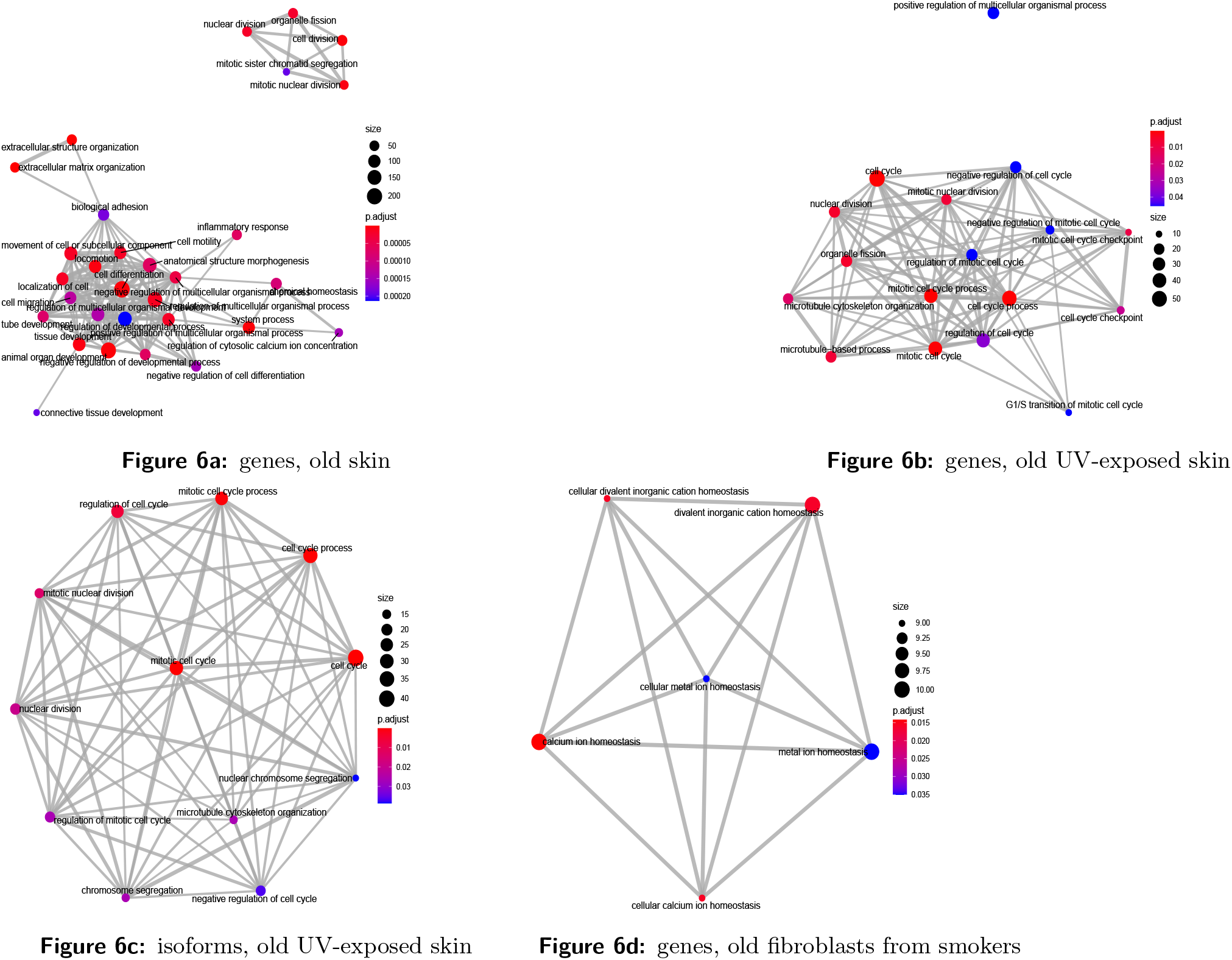
Enrichment maps of Gene Ontology Biological Process categories represented by significantly (q < 0.05) DE genes in old aged skin (a), genes (b) and isoforms (c) from male fibroblasts from UV-exposed skin, and old fibroblasts from smokers (d), identified using Salmon

GO:BP enrichment of genes and isoforms DE in old aged male skin identified genes involved in keratinization, keratinocyte and epidermal cell differentiation as well as skin development have higher expression levels in middle aged males. Enrichment maps of GO:BP terms identify reduced mitosis, extracellular matrix organisation, adhesion, cell differentiation, cell motility, calcium regulation alongside increased inflammation 6Figure 6a. These changes were mirrored by significant (q < 0.05) reductions in biological process categories relating to tissue/skin development, cell and keratinocyte, epidermal and epithelial differentiation, these processes were highly significant (q < 0.01) and part of cornification. Both genes and isoforms DE in fibroblasts from UV exposed skin identified reduced cell cycle, microtubule, cytoskeleton organisation and DNA maintenance 6Figure 6b, 6Figure 6c. In young vs old comparisons of fibroblasts from smokers the most significant processes related to calcium homeostasis 6Figure 6d. Functional enrichment of DM CpGs for genes identified genes involved in axon development, genesis and guidance are changed from middle to old age. Significant pathways are involved in related processes (axon guidance, regulation of the actin cytoskeleton), as well as Calcium signalling and related processes (cell adhesion, Notch, WNT, Rap1, platelet and phospholipase D signalling), further represented by cell-substrate adhesion/organisation, cell junction assembly/organisation and proteoglycans in cancer. Enrichment analysis for DM CpGs identified that in young vs old age group comparisons conserved non-coding elements identify interconnected genes representing cytoskeletal dynamics; Microtubule actin crosslinking (MACF1), cell adhesion; Fibronectin (FN1), Tenascin (TNR), integrin (ITGA), bone (BMP7, bone morphogenic protein), cell signalling serum response factor (SRF, regulates cell proliferation and differentiation), the calcineurin (Secreted Frizzled Related Protein, SFRP2), and the hormone responsive nuclear receptor (NR2E1). This was further supported by GO reduction plots showing nuclear division, organ development and extracellular matrix organisation are highly significant (q < 0.005).

## Methods

### Identifying transcriptomic studies for inclusion

RNA-seq data deposited in public repositories was searched for the terms age, sex/gender and fibroblasts to identify suitable studies for inclusion in this work. Three studies were identified (Fleischer et al. 2018^11^, Kaisers et al. 2017^9^, and Jung et al. 2018), key differences in the study designs are summarised in 12.

Table 12 identifies that fibroblasts were cultured using different methods and harvested after a different number of passages. Of particular concern was the Jung data set, which did not detail age, sampled males and females and cultured fibroblasts with fibroblast growth factor (FGF). The Jung data set was dropped from the analysis because it was not possible to determine whether differences were due to sex, disease status, bodily location of tissues used to obtain cells or growth factors. Additionally RNA extraction and purification techniques, library preparation, sequencing technologies and read lengths all differed, as well as the age and gender distributions of the samples assessed. Existing data was extended to include paired RNAseq data from fibroblasts, UV and non-UV exposed skin derived from the GTEx study. Samples were further stratified by race and ethnicity so that only Europeans were considered. The smoking status of male Europeans identified that enough samples were available (> 20 per group) to assess the impact of smoking on the transcriptome of aging skin and fibroblasts.

**Table 12.**
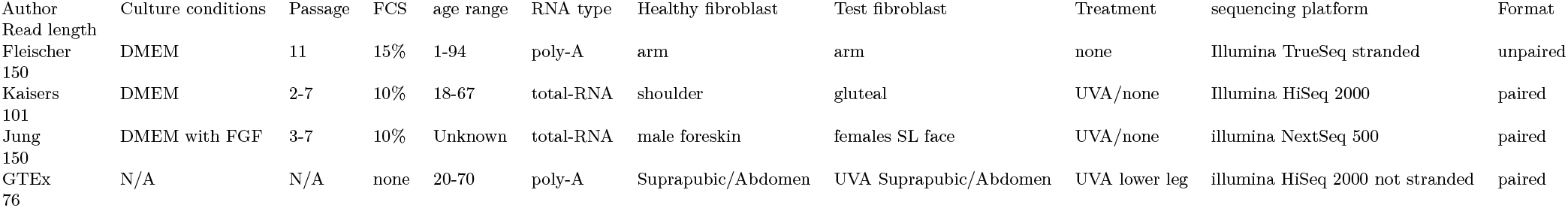
Summary of key differences in experimental conditions for each of the RNA-seq studies conducted on aged fibroblasts. Abbreviations FBS=Fetal Bovine Serum, FGF= Fibroblast Growth Factor

### Quality and distribution assessment

Quality was assessed using FastQC, quality, clustering and distribution was checked using Squared Cumulative Variance (CV^2^) plots using cummeRbund^38^, PCA plots and fitdistrplus^39^, which determined data followed a negative binomial distribution. Preliminary analysis and clustering identified sex specific expression of genes and isoforms, and more variable expression profiles in females, which strengthened the case for sex stratification. The final analysis methods were tweaked based on the availability of environmental exposure information relevant to skin ageing for GTEX samples and observed clustering.

### Derived GTEx data analysis methods

To maximise the likelihood of identifying DE genes and isoforms GTEx data was assessed using reference (HG38) based analysis with Salmon and CuffDiff, as well as using a *de novo* approach capable of identifying and quantifying novel transcripts.

### Reference based transcriptome quantification: Salmon

Alignment free quantification was completed on paired end fastq files to produce count tables which were imported and annotated using tximport^40^ with the gencode v32 gene map, subset into male and female age groups based on phenotypic data, normalised and analysed with DESeq2^41^, PCA plots identified that fibroblasts and skin were transcriptomically distinct. Samples were subsequently stratified by cell/tissue type, additional stratification by smoking and UV-exposure increased the number of DE genes identified as well as enrichment categories Network enrichment analysis on DE genes and transcripts.

### Reference based transcriptome quantification:CuffDiff

Paired end sequencing files were quality checked using fastqc, aligned to the HG38 genome using HISAT2, with the very sensitive setting, unpaired reads were excluded. SAM files were converted to sorted BAM files using SAMtools^23^ and insert sizes, mean inner distance, and standard deviation per sample was estimated using Picard tools (BroadInstitute 2014). Mapped reads were assembled into transcript files per sample using StringTie2^42^. FPKM normalised data was significance tested using the cross replicate pooled condition dispersion method in CuffDiff. CuffDiff was completed separately male age group comparisons, with and without UV-exposure and smoking status stratification. A dispersion model based on each replicated condition was built then averaged to provide a single global model for all conditions in the experiment. CuffDiff p-values were adjusted using Benjamini-Hochberg Multiple Testing Correction (BH-MTC)^43^ to generate q-values^44^.

### Network enrichment analysis on DE genes and transcripts

Significantly (q < 0.05) DE genes and transcripts from each of the stratification’s, comparisons and methods were assessed for functional and network enrichment using the R libraries clusterProfiler^45^, wordcloud^46^, org.HS.eg.db^47^, enrichplot^48^ and pathview^49^. Enriched gene ontology categories and pathways identified were plotted for visualisation using conserved non-coding elements plots, enrichment maps, go reduction plots and dot plots for the most significant pathways. Enrichment results were then separated according to where those changes occurred in cells or tissue layers and are presented in sections; ECM-fibroblast changes in the dermis Male skin: ECM and fibroblasts, cell/organelle changes in skin Male skin; cell and organelle level changes, epidermal tissue level changes Male skin: Epidermal changes.

### Methylation data selection

One publicly available data set (E-GEOD-5194) was selected for subsequent analysis, it contained mixed sex, mixed dermis and epidermis samples. Three dimensional PCA plots were generated using plotly^50^, ggplot2^51^, grid^52^, gridExtra^53^ and pca3d^54^ for each individual data set with age, sex, tissue type interpretations to identify factors influencing the clustering of samples; this identified sex and age group clustering for study E-GEOD-51954.

### Methylation data processing

Male idats were read using the read.metharray.exp function, all samples had detection p-values < 0.05 reported by the detectionP function in minfi and were retained for analysis, and annotation generated using the getAnnotation function in minfi^55^. Data was quantile normalised using male specific normalisation respectively, and probes with a detection p-value < 0.01 were discarded. Limma^56^ was used to generate model matrices and contrasts for age group comparisons (young < 30, Middle 30-65, Old > 65), empirical bayes was applied to the fit object for variance shrinkage, and Benjamini Hotchberg multiple testing correction applied to results, DM CpGs were determined if q-values were < 0.05.

### Functional enrichment of DM genes

Ensembl and Entrez identifiers for significantly (q < 0.05) DM CpGs were obtained from the *homo sapiens* database (org.Hs.eg.db)^47^ and enrichment determined using the R libraries clusterProfiler^45^, wordcloud^46^, and enrichplot^48^. Significant (q < 0.05) enrichment of Gene Ontology (Biological Process, Molecular Function, Cellular Component) terms and KEGG Pathways, were used to generate conserved non coding element, go reduction plots and enrichment maps from significantly (q < 0.05) DM CpGs associated with genes.

### Merging epigenetic and transcriptomic data: reference based CuffDiff

Annotated significantly (q < 0.05) DE genes and transcripts for male age group comparisons Derived GTEx data analysis methods, and annotated significantly (q < 0.05) differentially methylated genes Functional enrichment of DM genes were merged by nearest gene. Significantly DE transcripts that overlapped with DM CpGs were subset and are highlighted by an asterisk in figures.

## Discussion

### Cultured fibroblasts

In this study the impacts of ageing on gene and isoform expression in skin and fibroblasts were assessed using two DE quantification tools; Salmon, and CuffDiff. Initial analysis of fibroblasts focused on the larger, better annotated publicly available data sets (Fleischer et al. (2018)^11^ and Kaisers et al. (2017)^9^, analysed independently of one another. Creating age group categories and stratifying by sex reduced variance, increased sample sizes, and created data sets following a negative binomial distribution. The GTEx repository contained both fibroblast and skin samples, detailed age, sex, UV-exposure status and smoking status of participants, and covered a greater age range which made it highly advantageous, given the impact that these factors can have on skin ageing^1, 3, 4, 6, 7^. Clustering identified that fibroblasts were transcriptomically distinct from skin samples, so skin and fibroblasts were analysed independently. Smoking and UV-exposure; known to accelerate visible signs of skin ageing were used to generate additional stratification’s, hone in on ageing signatures, and investigate the impacts of these environmental factors. For fibroblasts results were enhanced by stratifying by smoking status (Figure 7), whereas in skin UV-exposure improved the results (Figure 9).

**Figure 7.**
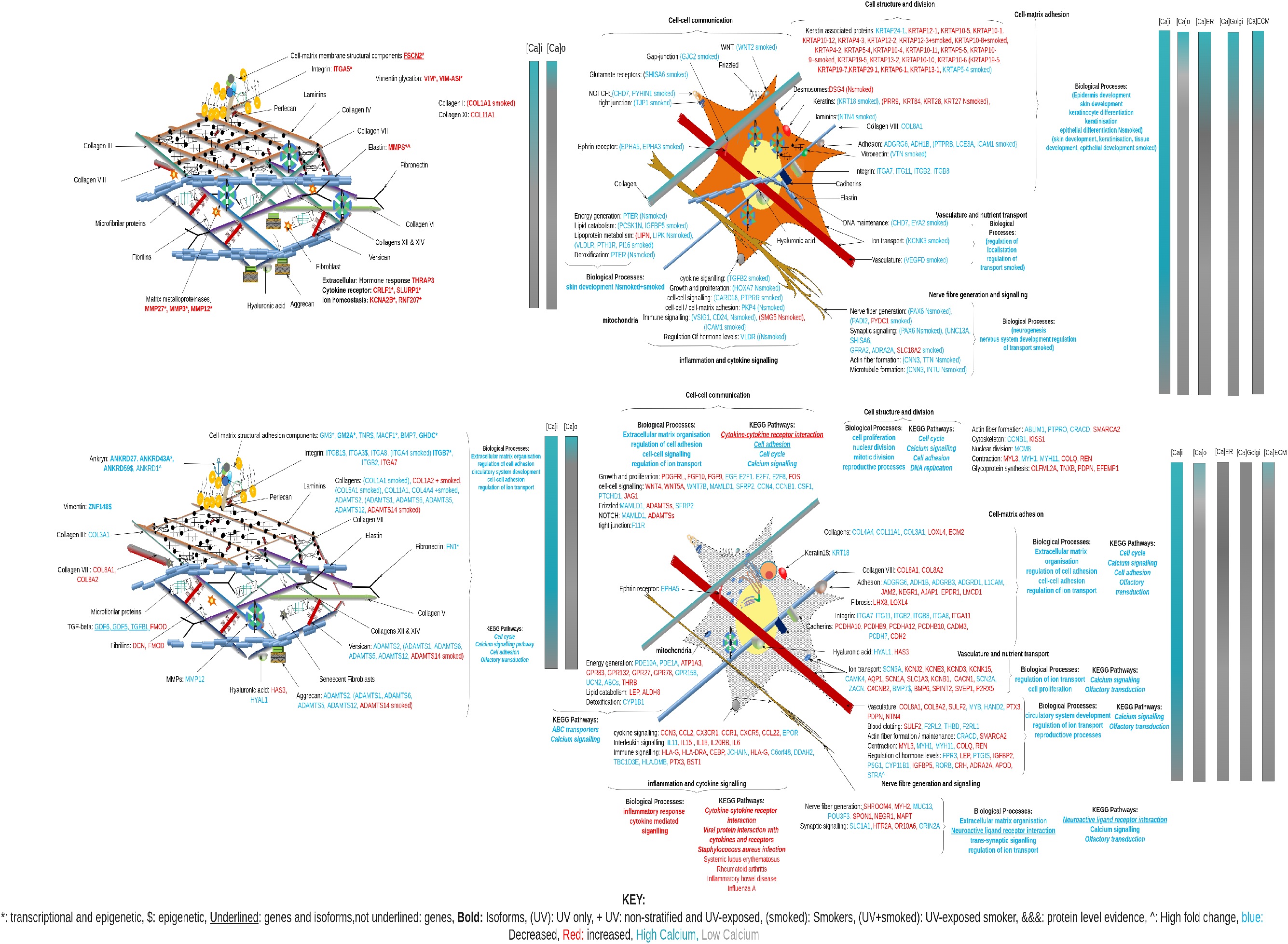
Molecular diagram of ECM and ECM-fibroblast interactions overlaid with transcriptomic and epigenetic changes for genes DE and DM, enriched GO Biological Processes and KEGG pathways for Middle age (top) and old age (bottom) ECM and fibroblasts, showing molecular consequences of those changes at a ECM and fibroblast level, supported by references^1, 2, 4, 7, 9, 13, 15–18, 20, 23–25, 30, 31, 33^

**Figure 8.**
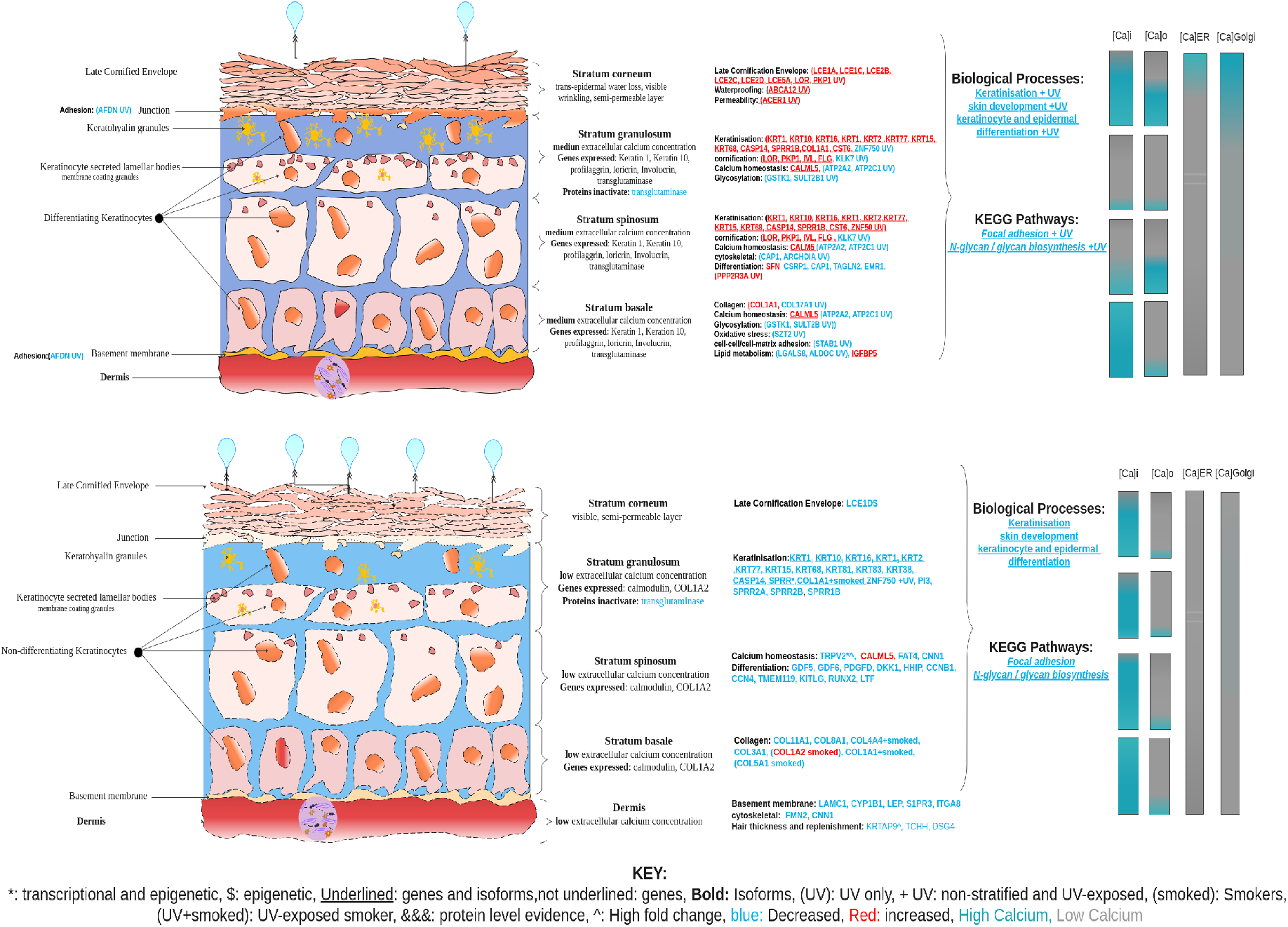
Molecular diagramatic overview of transcriptomic and epigenetic changes for genes DE and DM, and enriched GO Biological Processes and KEGG pathways for Middle age (top) and old age (bottom) keratinocytes, showing molecular consequences of those changes at a cell and organelle level. Supported by references^**?**, 2–4, 7, 13–17, 19, 19, 19, 22, 22, 26–29, 59–61^

**Figure 9.**
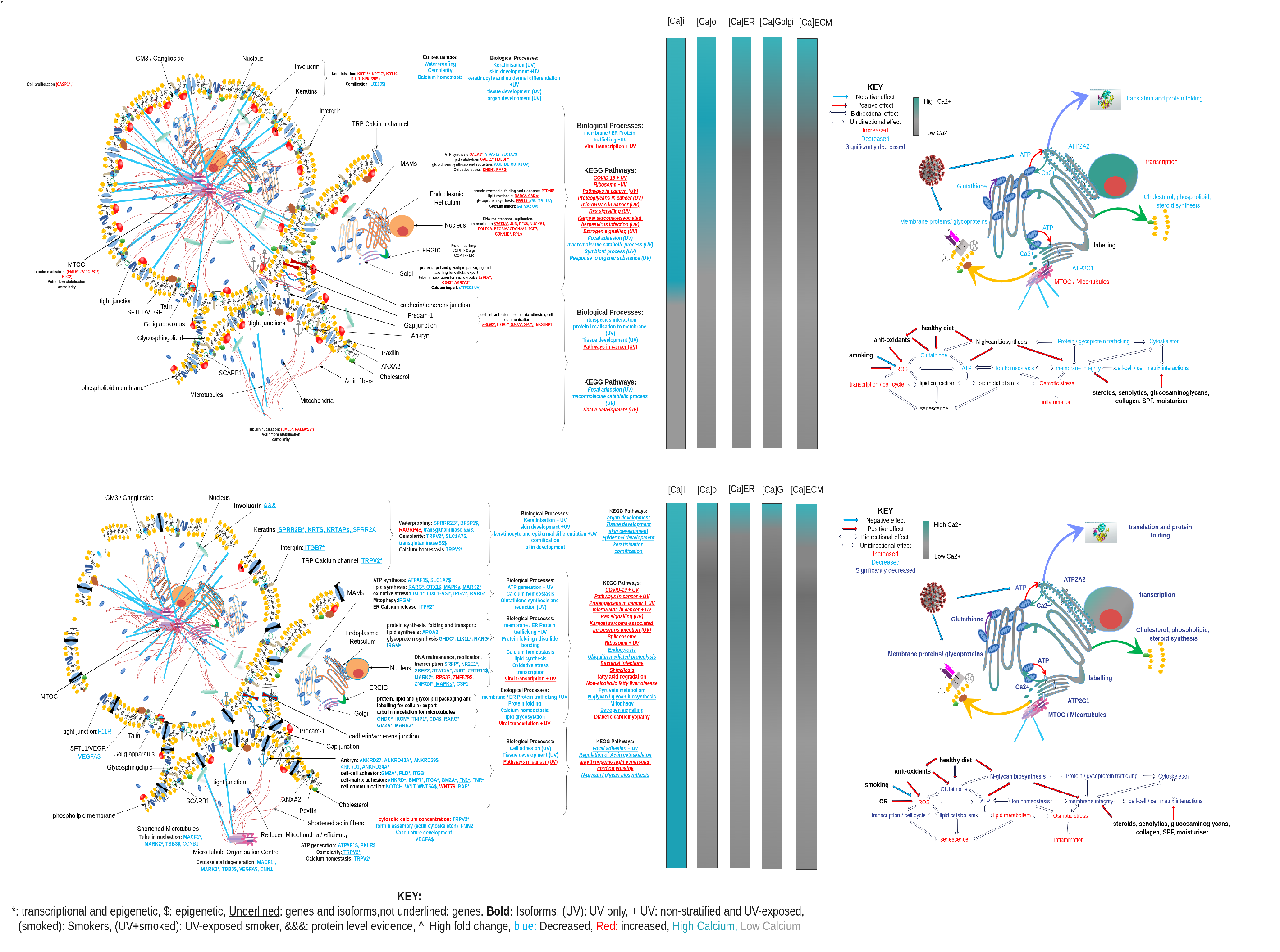
Diagram of the epidermis showing changes in the four layers and key features (calcium concentrations, genes expressed, and proteins activated) which facilitate the differentiation of keratinocytes and a depiction of the cellular and tissue wide consequences of ageing (middle age = top, old age = bottom) supported by references^1–4, 7, 9, 14–21^

### Male skin: ECM and fibroblasts

Functional enrichment (Gene Ontology, KEGG pathways) of genes and isoforms DE with age in fibroblasts identified alterations in cell adhesion, cytokine mediated signalling, vasculature development, synaptic signalling, and altered immune responses, regulation of hormone levels, G-protein signalling, ion transport, extracellular matrix organisation and skin development in old aged fibroblasts. Pathways identified alterations in infections, auto-immune disorders, cell adhesion and ABC transporters (Figure 7). There is evidence that culturing fibroblasts, particularly in the presence of fibroblast growth factor, increases fibroblast proliferation, alters the transcriptome, and the uptake of Ca^2+^ by mitochondria which can induce senescence^4, 7, 13, 21, 22^. Fibroblasts interact with and maintain the structural integrity of the ECM, which is known to be affected by ageing^1, 1–9, 13, 15–18, 23^. Extension of the study to GTEx fibroblasts identified affected genes and processes in GTEx fibroblasts and ECM 7 confirming increase expression of MMPs in middle age, implicating associated alterations in the organisation and structural integrity of collagen and elastin, reduced cell adhesion, as well as increased cytokine signalling and inflammation in old age (Figure 7).

### Ageing skin

In skin the differentiation of keratinocytes, and their waterproofing are influenced by extracellular Ca^2+^ concentrations, which can be affected by growth media constituents^21^, cell type^21, 22^, culture time^4, 13, 22^, and growth factor presence^7^; creating a potentially complicated set of confounding factors. Cultured human epidermal keratinocytes are known to increase the expression of cadherins, integrin, alpha-catenin, beta-catenin, plakoglobin, viniculin and alpha-actinin in response to increases in extracellular Ca^2+^^21^. This study assesses RNA-seq from uncultured whole skin, encompassing keratinocytes in the epidermis, the ECM and fibroblasts in the dermis. To aid interpret-ability of the complicated layers of information generated by the study, data are subset into cell and organelle Male skin; cell and organelle level changes, and the consequences of those changes in the formation and maintenance of epidermal tissue Male skin: Epidermal changes and presented diagrammatically in relevant sections.

### Male skin; cell and organelle level changes

In middle aged male skin genes increased in expression are involved in keratinisation, and cornification, unfolded protein responses; the functions of genes and isoforms increased in expression are associated with highly DM CpGs. They reveal repeated links to the MAM-ER-Golgi-ERGIC-cytoskeletal module 3, affecting Ca^2+^ handling, protein folding (PFDN5), lipid catabolism (PRR13, THBS1, HDLBP, GM2A, RARG), cytoskeleton formation and maintenance (cell cycle/proliferation CDKN1B, BTG2, JUN, THRAP3, NUCKS1), cell adhesion (FSCN2), and differentiation (SP7, FSCN2). The gangliosidase activator GM2A was increased in middle aged skin; dysregulation of GM2A has been associated with Gaucher disease; accumulation of glucosylceramide, with skin browning and peeling^22^. It irreversibly degrades gangliosides making them available as a substrate, and stimulating the release of intracellular Ca^2+^^27^. Rises in intracellular Ca^2+^ regulate keratinocyte differentiation pathways (keratinisation), via SPRR genes such as Small proline-rich protein 2B (SPRR2B, increased in expression and DM) leading to subsequent development of the cornified envelope^2, 16–19^. Cytosolic SPRR2B is cross-linked to the inner membrane by transglutaminase forming an insoluble barrier in the stratum spinosum^19, 29^. Pro-inflammatory signalling is thought to regulate SPRR and involucrin genes, which encourage wound healing, but are also implicated in inflammatory diseases and excessive growth conditions of the skin^19^. Evidence of unfolded proteins is provided by increased expression of cytosolic prefoldin alpha subunit (PFDN5, also known as MM1) it promotes the protection of actin and tubulin, playing an important role in cytoskeletal maintenance. It also enables the formation of beta-amyloid fibrils from oligomers (preventing the formation of beta-amyloid plaques seen in Alzheimer’s), alpha Synuelein fibrils from alpha synuclein oligomers (Parkinsons), and ensures the production of the non-toxic oligomer of HTT, reducing the production of inclusion bodies in Huntington’s disease^28^. Impairment of Ca^2+^ homeostasis in the ER leads to the accumulation of unfolded proteins, which if prolonged can lead to the activation of apoptosis^14^.

Interestingly in old age (> 65) cornification significantly reduced, consequently genes involved in keratinisation (KRTs, KRTAPs, SPRRs) and collagen (COL11A1, COL3A1, COL4A4, COL8A1) generation are decreased, leading to the identification of skin development functional categories 8. Functional enrichment of DM CpGs support these findings, identifying alterations in calcium signalling, phospholipase D, regulation of the actin cytoskeleton/axon guidance/actin filament organisation, cell adhesion and pathways implicated in cancer (WNT/NOTCH). Deregulation of calcium homeostasis is further supported genes with DM CpGs in old age, which included calcium signalling and related processes; regulation of the actin cytoskeleton/axon guidance/actin filament organisation, cell adhesion, Notch, WNT, Rap1, platelet and phospholipase D signalling) and DE genes in old male skin 6Figure 6a and fibroblasts from smokers 6Figure 6d. These decreases coincides with a decrease in the expression and DM of the promoter associated with cation-permeable Transient Receptor Potential calcium channel (TRPV2); a cation channel and regulator of PI3K/MAPK signalling^57, 58^. It’s decrease is seen alongside decreased expression and DM of promoters associated with the PI3K and MAPK signalling pathways; STAT5 (transcriptional and kinase activation) JUN (transcription factor, oncogene), MARK2 (Serine/Threonine protein kinase); dysregulation of which are implicated in cancer and ageing. These isoforms coincided with significant activation of pathways relating to renal cell carcinoma, proteoglycans in cancer, viral and bacterial infections (COVID-19, Shigellosis, *E*.*coli*, bacterial invasion of epithellia), cardiomyopathy and estrogen signalling.

COVID-19 infection is more severe in elderly males^59, 60^, this has been associated with ER stress resulting in an unfolded protein response, preventing protein trafficking, which dampens the immune response of the host, leading to inflammation, autophagy, and apoptosis of infected cells^61^, also see in Hepatitis C and cytomegalovirus^15^. The identification of viral pathways; COVID-19, KSHV, may therefore be due to overlapping molecular similarities; increased ER stress, unfolded protein response and autophagy, which are known features of ageing and viral infection^15, 22, 61–63^. TRPV2 and the related gene TRPV1 are involved in odorant detection and have been implicated in loss of smell symptoms and outcomes in COVID-19^64^, they are regulated by senolytics such as Fisetin found in spices, fruits and vegetables. Fisetin lodges in cell membranes, regulating Nrf2 antioxidant signalling pathways, upregulating expression and reduction of glutathione, preventing oxidative damage to lipids and glycation that block PI3k/AKT/mTOR pathways and mitosis^64–67^. High intakes of spices containing senolytics correlate with decreased COVID-19 mortality, improvements in non-alcoholic fatty liver disease (Figure 8) by activating Peroxisome Proliferator Activated Receptor PPAR signaling (RARG Figure 8)^68^, restoring SERCA expression improves sensitivity to insulin by addressing imbalances of Ca^2+^ in cardiac dysfunction in diabetes (diabetic cardiomyopathy 8) and reduced the presence of free fatty acids^**?**, 15, 64, 68, 69^. Specific channels sensitive to hormones and cytokines line cell membranes and are highly coordinated controlling the amplitude and duration of cytosolic Ca^2+^ spikes^24, 25^. The cytokine storm is a known, and dangerous complication of COVID-19 infection, leading to increased mortality^64^; cytokine signalling reduces calcium uptake from blood and extra-cellular fluids^70^, and thus altered cytokine signalling may occur to counter increased cytosolic Ca^2+^ which activates Protein Kinase C (PKC), calcineurin, calpains, calmodulin dependent kinases and calmodulin binding proteins, accelerated rates of NADH production and oxidative phosphorylation^23, 70^. Calcium stress inhibits mRNA translation to reduce the influx of new proteins into the ER, and activation of MAPK signalling^15, 16^; thus calcium dysregulation may also underpin metabolic changes. In ageing cells alterations in the cytokine signalling pathway are frequently observed alongside inflammation, and this has been linked to increased rates of senescence and cancer^23, 71^ also seen in old cells 8. Increasing the anti-oxidant potential of glutathione inhibits hormone like pro-inflamatory cytokines including TNF, ILK6 and NFK-B regulators of the immune system (such as TNIP1, GHDC, IRGM 8) and inflammation^72^. Proteins sensitive to Ca^2+^ transduce key signalling pathways which allows cells to respond appropriately to a range of stimuli such as depolarisation of cell membranes, osmolarity, cell surface distortion and changes in temperature^24^. It is therefore possible that aged individuals, particularly males, may have a more severe response to COVID-19 infection because infection compounds these pre-existing age related cellular stresses making the cells more susceptible to apoptosis.

Cellular calcium homeostasis is regulated by sex hormones 8, in males testosterone begins reducing after the age of 63^73^ and references therein. Interplay between sex hormones and calcium regulate cholesterol levels and *de novo* lipid synthesis determining whether a cell can undergo mitosis (^73^ supplementary discussion pages 3-5). Hormones have been implicated in regulating oxidative stress, lipid profiles; both testosterone and oestrogen have both been implicated in regulating lipid profiles, and increasing the activity of the plasma membrane calcium pump (TRPC)^74–78^.

The most significantly activated pathways were age related degenerative diseases; Alzheimer’s disease, amyotrophic lateral sclerosis, Huntington disease, Parkinson’s disease. Age related degenerative disorders including Alzheimer’s, lateral sclerosis, type 2 diabetes, obesity, GM1 gangliosidosis and viral infections including cytomegalovirus and hepatitis C have been associated with age related declines in the function of Mitochondrial Associated Membranes (MAM)^22^, associated Ca^2+^ dysregulation^15^, aberrant metabolism, decreased lifespan, increased ROS^22^, and disruption of the mitochondrial surface calcium channel ITPR2; an inducer of cellular senescence^26^. MAM are involved in processes such as lipid biosynthesis and trafficking, reactive oxygen species (ROS) production, autophagy^22^ expression of ER proteins and the formation of disulfide bonds^15^; their disruption is supported by middle aged increases in PFDN5, RARG and GM2A, followed by reductions in high density lipid catabolism (RARG, GM2A, LIX1L, TNIP1, GHDC and IRGM) in old age. MAMs also regulate mitochondrial fission and fusion, essential for mitochondrial quality control; in ageing mitochondrial fusion increases with age, and fission decreases (Fis1)^22, 26^, in this study genes involved in mitochondrial fission were decreased in old age. Providing evidence of decreases in mitochondrial numbers, regeneration, ATP production, energy declines and associated reductions in high density lipid catabolism and transport within and between cells^22^.

Given the structural role of calcium, in terms of stabilisation of disulfide bonds, formation and stabilisation of the cytoskeleton, and adhesion of cells^15, 20, 24, 25, 30, 33^ Introduction. Calcium dysregulation could also explain decreases in expression and DM of CpGs associated with calcium binding (SFRP2), cytoskeletal development (Microtubule actin crosslinking factor 1 MACF1), cell adhesion (Integrin subunit alpha 3 ITGA3, Fibronectin FN1, Tenascin TNR, bone morphogenic protein (BMP7), Ankyrin (ANKRD1). Which are constituents and active members of the affected neuroactive ligand-receptor interaction, ECM-receptor interaction pathways, focal adhesion, and regulation of actin cytoskeleton, represented by age related decreases in cytoskeletal formation in ageing cells 8. A further consequence of intracellular Ca^2+^ alterations is reduced production of cytoskeletal projections required for mitosis (nuclear division), axonogenesis, and organelle fission leading to reduced cell adhesion, differentiation, pushing the cells (keratinocytes and fibroblasts) into a senescent state with decreased synthesis of the extracellular matrix.

### Male skin: Epidermal changes

For skin significant DE was identified for male age group comparisons, with most DE genes identified in young vs middle age group comparisons (i.e. the target age group where skin ageing becomes visible). Genes and isoforms identified using both CuffDiff and Salmon, and DM CpGs identified using limma were functionally enriched 8.

When samples were stratified by UV-exposure status CuffDiff isoforms had higher expression of LOR (loricin), filaggrin (FLG), involucrin (IVL), Collagen I (COL1), the Late Cornification Envelope (LCE), Keratins 1 and 10, (KRT1 and KRT10), SPRR1B and CALML5 in middle aged UV-exposed male skin 5Figure 5c. Expression of these genes is regulated by the calcium switch (calmodulin, CALML5), which triggers expression of involucrin, loricrin, transglutaminase, Keratins (1 and 10), fillaggrin, to form cornified envelopes^16^. This process is tightly linked to a rise in intracellular free calcium, which is raised in response to increases in extracellular calcium concentrations^16^. UV-exposure is known to increase 7-dehydroxycholesterol (pre-cursor for vitamin D) production, alter the expression of SPRR genes, in conjunction with activating stress and a differentiation pathways such as calcium signalling, which regulate gene/protein expression and development of the late cornified envelope^2, 16–19^, all of these changes were seen in middle aged skin 9. Alterations in calcium homeostasis are further supported by Salmon which identified lower expression of the Golgi calcium pump (ATP2C1) and the ER calcium pump (ATP2A2) 5Figure 5b in UV exposed middle aged male skin. Both ATP2C1 and ATP2A2 require a supply of ATP from the mitochondria, which in association with MAMs produce the antioxidant GSTK1 (glutathione s-transferase) decreased in middle aged males. Activity of SERCA Ca^2+^ import pumps located on the ER membrane is regulated by N-glycosylation, glutathionylation, and calcium/calmodulin kinase-II dependent phosphorylation, as well as in response to phospholamban (PLB) and sarcolipin (SLN),^14^, and these changes are associated with altered expression of calmodulin (CALM5) 5Figure 5c, 9. Decreased expression of these genes is evidence of reduced Ca^2+^ uptake into the ER and golgi apparatus, associated with increased intracellular Ca^2+^, leading to reduced cell adhesion, increased K10 expression, and an overall reduction in keratinisation^2, 16–19^. Consequently significant Gene Ontology categories represent alterations in keratinisation, keratinocyte differentiation, protein targeting to the membrane and ER, cell surface receptor signalling, biological and cell adhesion 9. All of these changes are seen in middle aged UV-exposed male skin, indicating disruption of the membrane barrier and disruption to the calcium gradient. Waterproofing, and cornification steps are dependent on transglutaminase activity, which is regulated by intracellular Ca^2+^, which is in turn regulated by extracellular Ca^2+^ and thus disruption of the calcium gradient, disrupts cornification^2^, which could explain age related increases in trans-epidermal water loss^6^ and observed dermal thinning^3, 7, 9^.

In old age (> 65) cornification is the most significantly reduced process, consequently genes involved in keratinisation (KRTs, KRTAPs, SPRRs) and collagen (COL11A1, COL3A1, COL4A4, COL8A1) generation are decreased, leading to the identification of skin development functional categories 9. In old aged male skin genes involved in keratinocyte development, differentiation, and adhesion are significantly decreased in expression and DM, functional enrichment of DE and DM genes points to cation homeostasis; specifically reduced ER Ca^2+^ import by ATP requiring TRPV2. Since intracellular Ca^2+^ concentrations regulate lipid (RARG, LIX1L, GM2A) and glycoprotein synthesis (GHDC, IRGM, HDBP, TNIP1), cytoskeletal formation (MARK2), cell adhesion (ITGB7), and thus the structural integrity and osmotic potential of cells. Decreased cell cycle (RARG, STATS, MARK2, JUN) and cytoskeletal maintenance (MARK2), are evidence of cellular senescence, which may relate to ROS (LIX1L), or Ca^2+^ homeostasis (TRPV2), or any of the inter-related processes.

The results suggest that reductions in ER/golgi Ca^2+^ import seen in middle aged male skin led to increased intracellular Ca^2+^, which initially drove an increase in expression of genes involved in keratinisation and cornification processes. As a consequence of loss of ER and golgi Ca^2+^ import, decreased Ca^2+^ concentrations in stratum granulosum reduced the activity of transglutaminase^**?**^ which disrupted formation of the late cornification envelope (decreases in LCE1D_cg01406203), leading to age related increases in trans-epidermal water loss^6^ and dermal thinning^3, 7, 9^.

### common ageing signatures in skin and fibroblasts

In this study extension to skin confirmed age related alterations in Ca^2+^ homeostasis seen in cultured fibroblasts, identifying additional key changes in keratinisation and cornification which may underpin age related decreases in the hydration, collagen content and thickness of skin^6^. Skin development is regulated by an extracellular calcium gradient^16, 17^; disruption to calcium handling at a cellular level, can impede this gradient, deactivating transglutaminase, decreasing membrane cross-linking, leading to thinning, and reduction in the waterproofing of the late cornified envelope; linked to the skin disorders Dariers and Hailey-Hailey disease^16, 17^. Disruption of the late cornified envelope increases trans-epidermal water loss, linked to decreased Glycosaminoglycans (GAGs) such as chondroitin and hyaluronic acid, potentially explains age related increases in skin thinning and dryness^6^, which are known^4, 6^ and were shown to be exacerbated by UV-exposure in males in this study Male skin: Epidermal changes.

## Acknowledgements (not compulsory)

The authors would like to thank Dr Simon Cockell for his time, support, and valuable advice and expertise.

## Author contributions statement

J.C., J.W., and L.I.P analysed public fibroblast transcriptomic data. L.I.P., J.W. and D.S, re-designed the analysis, L.I.P. analysed the results, drew the diagrams and wrote the manuscript. All authors reviewed the manuscript.

## Additional information

To include, in this order: **Accession codes** (where applicable); **Competing interests** (mandatory statement). The corresponding author is responsible for submitting a competing interests statement on behalf of all authors of the paper. This statement must be included in the submitted article file.

